# Effectiveness of heat tolerance rice cultivars in preserving grain appearance quality under high temperatures - A meta-analysis

**DOI:** 10.1101/2023.03.09.531821

**Authors:** Hitomi Wakatsuki, Takahiro Takimoto, Yasushi Ishigooka, Motoki Nishimori, Mototaka Sakata, Naoya Saida, Kosuke Akagi, David Makowski, Toshihiro Hasegawa

## Abstract

**Background:** Climate change, particularly rising temperatures, negatively affects rice grain quality, increasing chalky grain percentage (CG) and hampering rice grade and price. Heat-tolerant cultivars have been bred and released since the 2000s, but the effectiveness of heat tolerance in reducing the occurrence of CG has yet to be quantified.

**Objectives:** This study aimed to measure the effectiveness of breeding for better heat tolerance in reducing the negative impact of high temperatures on rice quality.

**Methods:** Through a systematic literature search, we developed a dataset including 1297 field observations covering 48 cultivars from five different heat tolerant ranks (HTRs) at 44 sites across Japan. A linear mixed-effect model (LME) and a random forest model (RF) were fitted to the data to analyze the effect of HTR and climatic factors such as the cumulative mean air temperature above 26 °C (TaHD), mean solar radiation, and mean relative humidity for 20 days after heading on CG.

**Results:** The LME model explained 63 % of the variation with a 14% RMSE. The RF partial dependence plot revealed that the logit-transformed CG response to climate factors was linear, supporting the assumption of LME. The statistical analysis showed that CG increased as a function of TaHD (P < 0.001), with significant differences among HTRs (P < 0.001). The strongest effect of TaHD was obtained for the lowest HTR and was found to decrease with increasing HTR. CG also increased with higher relative humidity (P < 0.001) and solar radiation (P < 0.01). Based on our modeling, we estimated that as TaHD increased from 20 to 80 °Cd (equivalent to a mean temperature increase from 27 °C to 30 °C), CG increased by 66 % points (difference in CG) for cultivars with the lowest HTR, 45 % points for cultivars with an intermediate HTR, and 19 % points for cultivars with the highest HTR. Raising HTR by just one step (from intermediate to moderately tolerant) is projected to increase the proportion of first-grade rice at a grain-filling temperature of 27 °C, but tolerance levels need to be improved further in case of stronger warming.

**Conclusions:** The effect of high temperatures on CG was highly dependent on the cultivar’s HTR. Improvements in HTR effectively reduce the negative impacts of high temperatures on rice grain quality.

**Significance:** Heat-tolerant cultivars are projected to suppress the prevalence of CG more than threefold compared with heat-sensitive cultivars when grain-filling temperature increases from 27 to 30 °C.

## 1. Introduction

Ongoing climate change, particularly rising temperatures, negatively affects rice grain yield and quality (Krishnan et al.,2011). Since rice (Oryza sativa) is mostly consumed as whole grain, the quality of rice involves a few unique traits different from other cereals, including milling, eating and cooking properties, grain appearance, and nutritional quality (Neerja and Renu,2020). Among various quality traits, appearance quality, especially grain chalkiness, has been of primary concern in major rice-production areas in the world because it affects the milling, eating, and cooking properties and is strongly affected by climate change (Cheng et al.,2005; Fitzgerald et al.,2009; Chun et al.,2010; Lanning et al.,2011; Ali et al.,2019). Not only does the occurrence of chalky grain (CG) lower the market price, but milling losses caused by fragile CG also reduce head rice yield and create direct income reduction (Lyman et al.,2013). A few simulation studies estimated increasing economic losses due to the further degradation of grain quality resulting from increased temperatures under climate change by the mid-century in Japan (Masutomi et al.,2019) and in Europe (Cappelli and Bregaglio,2021).

The chalkiness of grain is due to the presence of an opaque part in the grain endosperm, and is often classified into milky-white, basal-white, and white-back/belly, depending on the locations where opaque parts appear(Nagato and Ebata,1965). It looks opaque because light reflects irregularly in endosperms with larger air spaces formed by loosely packed starch granules, often caused by insufficient starch accumulation induced by high temperatures during the grain-filling period (Morita,2008). The occurrence of CG increases as the temperature during the grain-filling period increases. In Japan, as extremely high temperatures have become more frequent since the beginning of the 1990s (Japan Meteorological Agency,2015), CG has been increasingly reported across the country by the Ministry of Agriculture, Forestry, and Fisheries of Japan (MAFF,2021). To reduce the heat-induced damage on appearance quality, genetic improvement of heat tolerance is considered effective, and heat-tolerant cultivars have been bred and released since the 2000s (Ishimaru et al.,2016). The cultivation area of heat-tolerant cultivars increased from 2.3 % in 2010 to 11 % in 2020, and it is expected to expand more (MAFF,2021).

However, heat tolerance levels differed even among ’heat-tolerant cultivars’. In 2018, MAFF established nine heat tolerance ranks (HTRs) to enhance heat tolerance breeding. In addition, assigned several check cultivars to each HTR as part of the Plant Variety Protection and Seed Act (No. 83 of 1998)(MAFF,1998; Kaji et al.,2016; Tamura et al.,2018). Rice breeders can now have a clear breeding target and determine the HTRs of new rice cultivars by comparing their proportions of CG in high-temperature settings (mean temperature for 20 days after heading of around 27 °C) with those of check cultivars. HTRs could potentially be used to provide quantitative levels of heat tolerance of rice cultivars at different temperature levels, but the effects of HTRs on CG are yet to be quantified. A lack of quantified effectiveness of different HTRs in response to temperatures limits our ability to develop quantitative breeding targets in specific regions and at specific global warming levels.Many experimental studies have investigated the effects of high temperatures on grain appearance quality of various cultivars across Japan for different purposes, including variety selection and management improvement (See a list of references in Supplementary Materials). A few studies have quantified the effects of air temperature and solar radiation on the occurrence of CG in specific conditions (Okada et al.,2011; Masutomi et al.,2015; Takimoto et al.,2019). Masutomi et al. (2022) conducted a parameter sensitivity test to demonstrate the importance of CG dependence on temperature in the impacts of climate change on rice grain quality. It remains unclear, however, how much actual cultivars with high HTRs can alleviate the negative impacts of ongoing and projected climate change, which is necessary to develop concrete breeding targets in different regions and global warming levels.

Previous modelling studies often used parameters for individual cultivars, referred to as “genetic coefficients” (e.g., Mavromatis et al.,2001), but lack of data availability frequently limits model applicability for a large number of cultivars. An option would be to use the same set of coefficients for all cultivars belonging to the same HTR groups, thus reducing the need for cultivar-specific coefficients. Quantifying the heat tolerance for HTRs has at least two advantages over quantifying it for each cultivar. First, the effectiveness of increasing heat tolerance levels can be easily quantified in response to global warming levels, leading to the development of breeding targets for specific areas and temperatures. Second, each parameter can be derived using more observations covering a wide range of environments, improving the robustness of parameters and estimates. To utilize HTR as a quantitative measure, we need to examine if cultivars in the same HTR behave similarly in response to major climatic variables and if high HTRs have clear advantages over lower HTRs with increasing temperature levels. To answer these questions, we need a large dataset covering cultivars with various HTRs obtained from widely different environments. In this study, we conducted a systematic literature and database search to develop a nationwide dataset on rice grain appearance quality of cultivars of different HTRs. We then performed a meta-analysis to model the effects of temperature and other weather conditions on the CG of various HTRs. The results are used to examine the effectiveness of cultivar improvement in reducing the negative impact of temperature on rice quality.

## 2. Material and Methods

### 2.1. Selection of studies

We conducted a systematic literature search from three different sources to obtain CG data of rice cultivars grown under field conditions (Figure 1).

**Figure 1.**
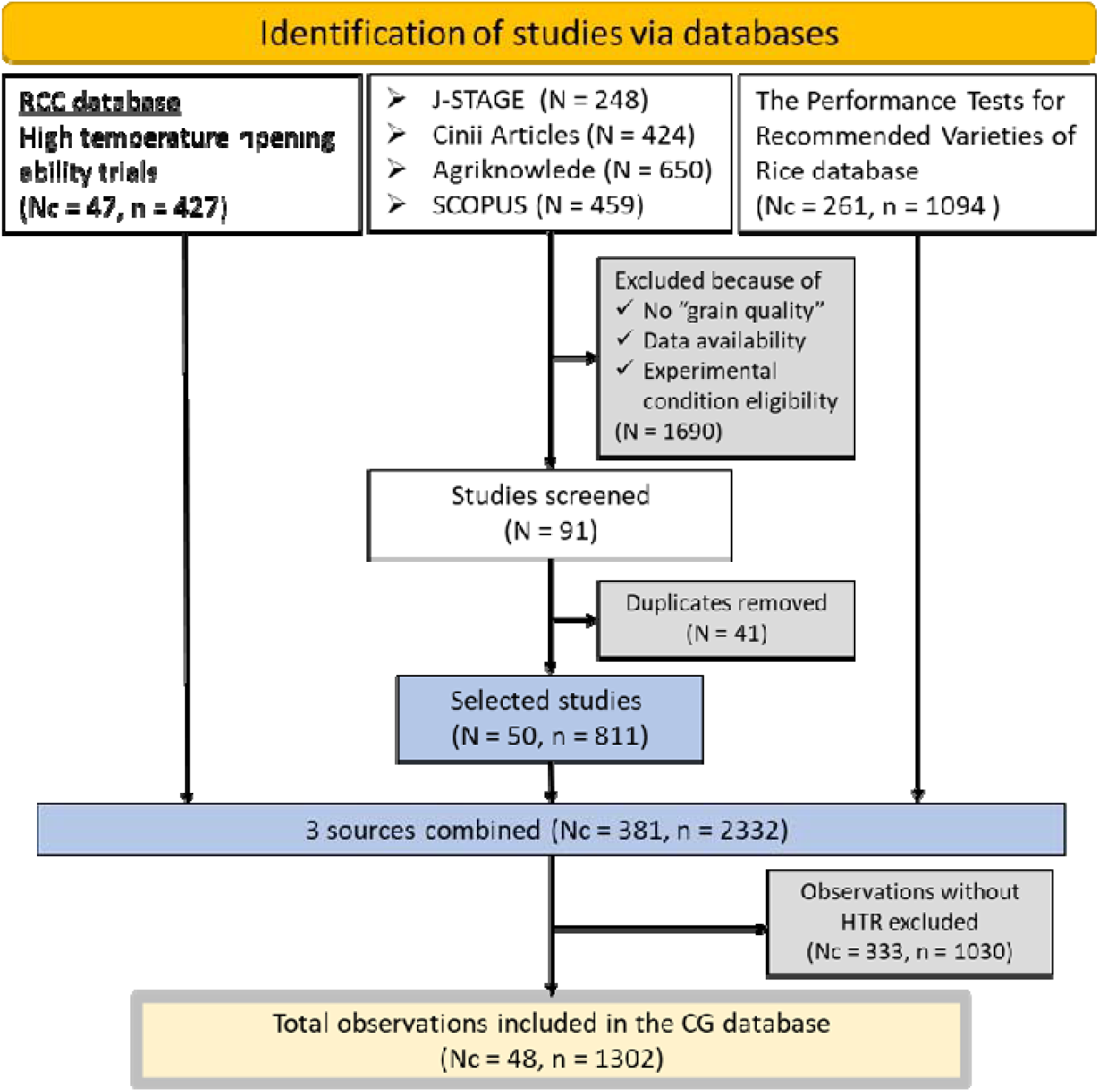
A PRISMA diagram depicting data collection and selection processes. Nc is the number of cultivars, N is the number of studies (articles) found through the literature search, and n is the number of observations.

The first data source is the Rice Cultivar and Characteristics (RCC) database developed by the Institute of Crop Science, NARO (https://ineweb.narcc.affrc.go.jp/), containing breeding progress, morphological and ecological characteristics, and appearance and taste quality characteristics of more than 500 varieties registered by MAFF. In this study, we focused on 63 cultivars: 39 cultivars selected as standard cultivars of different heat tolerance ranks under the Plant Variety Protection and Seed Act (No. 83 of 1998)(MAFF,1998), and 24 cultivars released as heat-tolerant cultivars (Supplementary Table1). Using 63 cultivars’ names as search terms, we searched experimental data in the RCC database and selected relevant data according to the following criteria. i) Experiments were conducted under open field conditions without warming treatments. Pot experiments and glasshouse experiments were excluded. ii) Experimental results should include CG, heading dates, and precise experimental field locations. In this study, we defined CG as the total % of milky-white, basal-white, belly-white, and back-white grains. We identified 427 observations of 32 cultivars to meet our criteria from the database.

The second data source consists of four databases containing peer-reviewed scientific papers and bulletins of national and prefectural agricultural research stations in Japan:

1. J-STAGE (https://www.jstage.jst.go.jp/browse/-char/en) ii) CiNii Research (https://cir.nii.ac.jp/?lang=en)
2. SCOPUS
3. AgriKnowledge, a database for agricultural science and technology of Japan (https://agriknowledge.affrc.go.jp/).

In December 2021, we searched J-STAGE and CiNii Research, using "Kouon-tojuku" in Japanese, meaning “grain filling under high temperatures”, as a search term, and obtained 248 and 424 results, respectively. For the SCOPUS search, the following equation was used;

**Search equation:** PUBYEAR > 2001 AND (TITLE-ABS((("heat stress tolerance" OR "heat tolerance" OR "heat stress") AND ("genotyp*" OR "cultivar" OR "grain quality" OR "nutrient quality"))) OR

AUTHKEY((("heat stress tolerance" OR "heat tolerance" OR "heat stress") AND ("genotyp*" OR "cultivar" OR "grain quality" OR "nutrient quality")))) AND (TITLE-ABS({greenhouse gas} OR {global warming} OR {climate change} OR {climatic change} OR {climate variability} OR {climate warming}) OR AUTHKEY({greenhouse gas} OR {global warming} OR {climate change} OR {climatic change} OR {climate variability} OR {climate warming})) AND NOT ((TITLE-ABS(MRV) OR AUTHKEY(MRV))) MRV stands for Measurement, Reporting, and Verification, the process for quantifying greenhouse gas emissions, and was used with “NOT” to exclude papers that focused on climate change mitigation. The SCOPUS search resulted in 459 articles. In June 2022, we conducted an additional search in AgriKnowledge using the 63 cultivars’ names as a search string and collected 650 articles. From a total of 1781 papers obtained from the four databases, we manually selected studies using the same two criteria described earlier. This manual selection screened 1691 papers and returned 90 studies. After removing duplicates, 50 studies (articles) were selected containing 811 field observations of 128 cultivars.

The third data source was the Performance Tests for Recommended Varieties of Rice data of Kochi Prefecture from 2013 to 2020, provided by the Kochi Prefectural Agricultural Research Center, containing 1094 sets of field observations of 261 cultivars. In all three sources combined, the dataset consists of 2332 observations of 381 cultivars collected between 1994 and 2020 in 46 experimental sites covering 33 prefectures (Figure 2).

**Figure 2.**
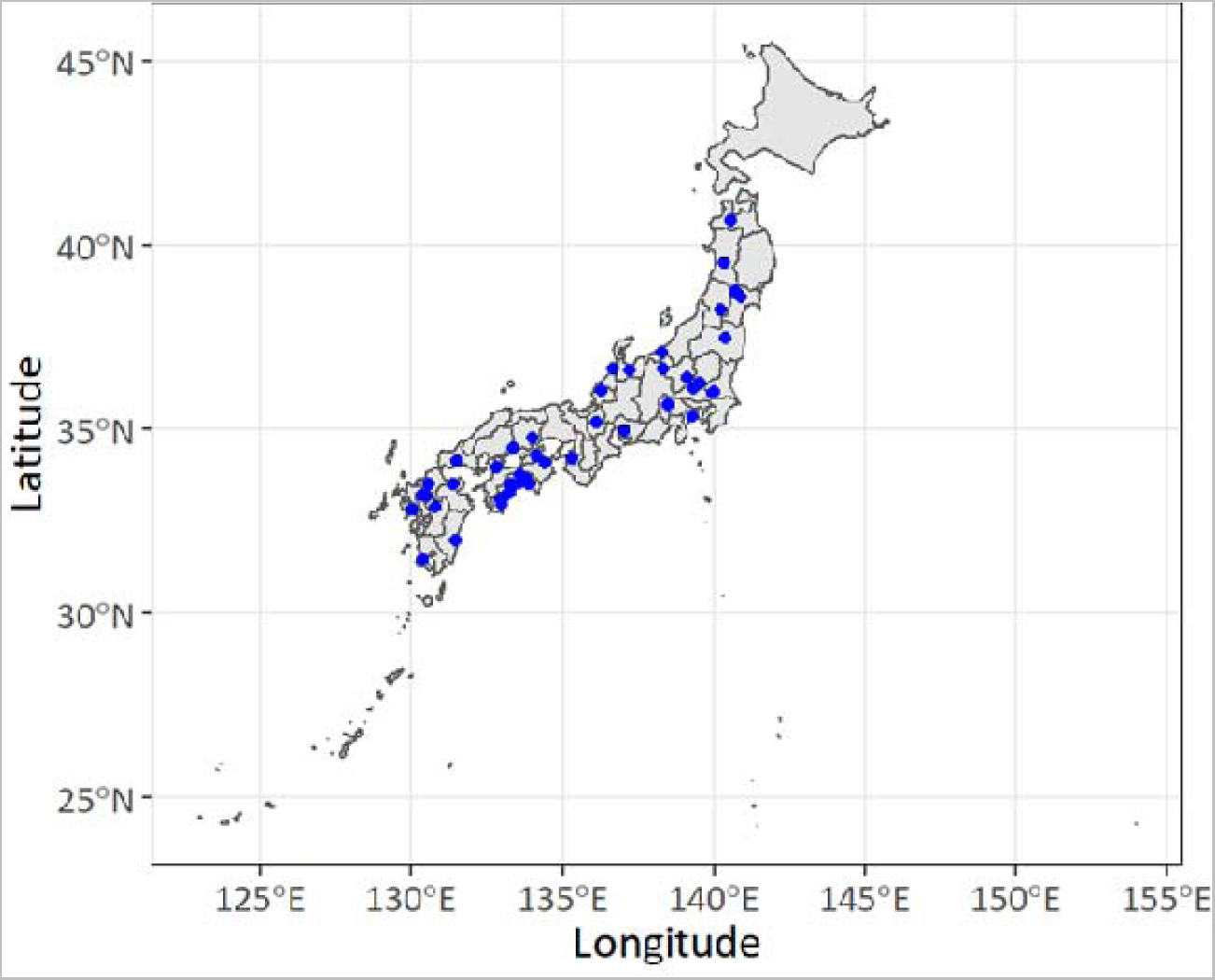
Geographical distribution of the 46 experimental sites in the rice quality dataset in Japan.

### 2.2 Data extraction

We extracted attributes such as prefecture, location of experimental sites, latitude and longitude (when available), soil group, cultivar name, heading date, and grain quality for 2332 observations (Table 1). Soil group to soil classification groups for each experimental site based on literature values or classified using the Soil Inventory website. (https://soil-inventory.rad.naro.go.jp/figure.html)(NARO). In addition to the variables listed in Table 1, the dataset contains sieve size for grain screening and CG measurement methods, such as “manual visual inspection” and “mechanical grain discriminator”. The standard errors and numbers of replicates were not available in the selected studies and could thus not be extracted. Of the 321 cultivars in our dataset, 48 have been classified in one of the existing heat tolerance ranks, while the rest have not been classified. Of the 48 cultivars with known HTRs (Table 2), 33 were listed in the MAFF guideline, ten were rated in reference to the standard cultivars, and their HTRs were reported in the RCC database. HTRs of the other five cultivars were obtained from the literature (Wada et al.,2010; NARO,2011; Mizobuchi et al.,2018; NARO,2018) (Table 2). A total of 1302 observations of the 48 classified cultivars were registered in our dataset. However, data in the zero nitrogen fertilizer plots (n = 5) were excluded from the following statistical analysis because nitrogen deficiency was reported to increase CG substantially (Wakamatsu et al., 2008) and hinders cultivar comparisons. The 48 cultivars registered in our dataset belonged to HTRs ranging from 3 (susceptible) to 7 (tolerant) (Table 2). Currently, cultivars with HTRs higher than 7 or lower than 3 have not been registered.

**Table 1.** List of variables obtained from the systematic literature search.

2.2 Data extraction
| Variables | Unit | Definition |
| --- | --- | --- |
| Prefecture |  | Prefecture of experimental site |
| Municipality |  | Municipality of experimental site |
| Institution/Station |  | Name of institution and or experimental station |
| Latitude | °N | Latitude of experimental site |
| Longitude | °E | Longitude of experimental site |
| Altitude | m | Altitude of experimental site |
| Soil group |  | Soil classification of experimental sites |
| Year |  | Year of experiment |
| Cultivar |  | Name of cultivar |
| Heading DOY |  | Heading day of year |
| HTR |  | Heat Tolerance Rank (HTR) from 3 (susceptible) to 7 (tolerant) |
| Tave | °C | Average temperature of 20 days after heading |
| TaHD | °C d | Cumulative mean air temperature above 26 °C for 20 days after heading |
| SR | MJ m<sup>-2</sup> d<sup>-1</sup> | Average solar radiation for 20 days after heading |
| RH | % | Average relative humidity for 20 days after heading |
| CG | % | % of chalky grains, a total % of milky-white, basal-white and belly/back-white grain |
| Milky-white | % | % of milky-white grain type CG |
| Basal-white | % | % of basal-white grain type CG |
| Belly/back-white | % | % of belly/back-white grain type CG |
| Reference |  | Source of the data records |

**Table 2.**
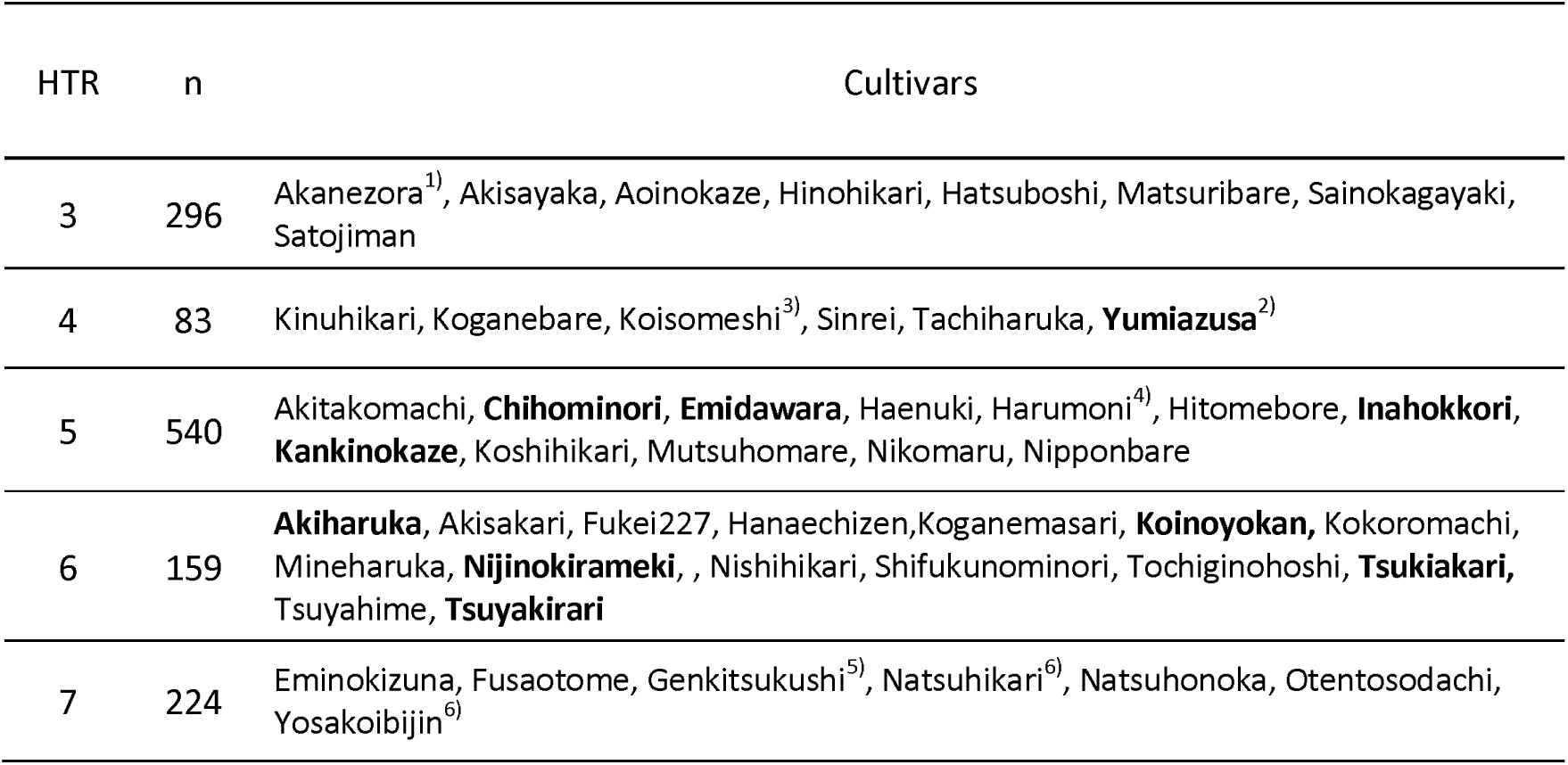
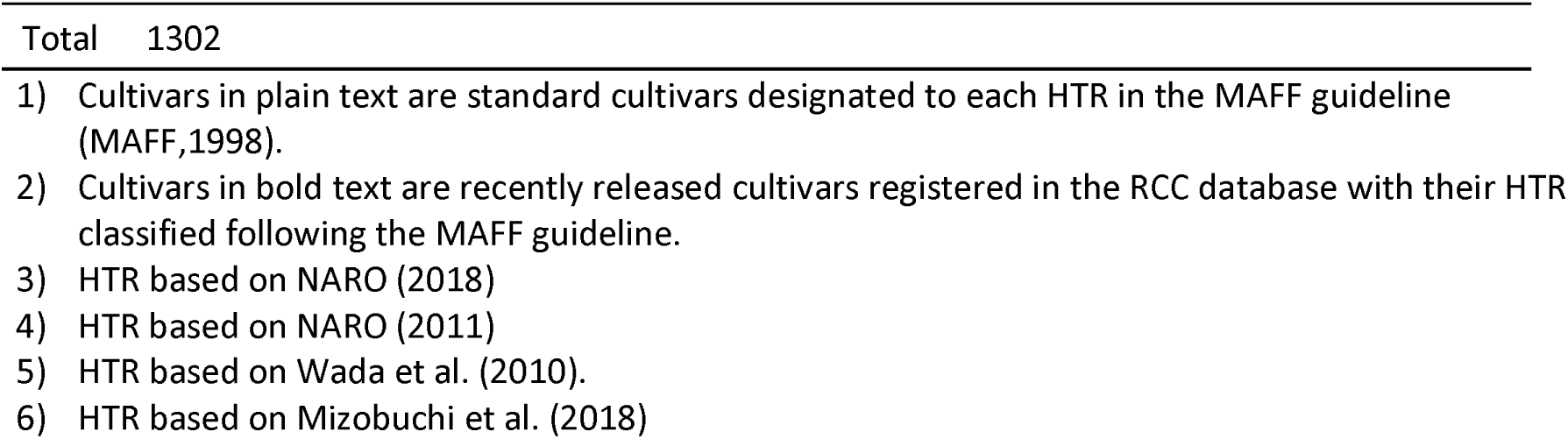
Number of observations (n) and names of cultivars for each heat tolerance rank (HTR) group.

| HTR | n | Cultivars |
| --- | --- | --- |
| 3 | 296 | Akanezora<sup>1)</sup>, Akisayaka, Aoinokaze, Hinohikari, Hatsuboshi, Maturibare, Sainokagayaki, Satojiman |
| 4 | 83 | Kinuhikari, Koganebare, Koisomeshi<sup>3)</sup>, Sinrei, Tachiharuka, **Yumiazusa**<sup>2)</sup> |
| 5 | 540 | Akitakomachi, **Chihominori**, **Emidawara**, Haenuki, Harumoni<sup>4)</sup>, Hitomebore, **Inahokkori**, **Kankinokaze**, Koshihikari, Mutsuhome, Nikomaru, Nipponbare |
| 6 | 159 | **Akiharuka**, Akisakari, Fukei227, Hanaechizen, Koganemasari, **Koinoyokan**, Kokoromachi, Mineharuka, **Nijinokirameki**, Nishihikari, Shifukunominori, Tochiginohoshi, **Tsukiakari**, Tsuyahime, **Tsuyakirari** |
| 7 | 224 | Eminokizuna, Fusaotome, Genkitsukushi<sup>5)</sup>, Natsuhikari<sup>6)</sup>, Natsuhonoka, Otentosodachi, Yosakoibijin<sup>6)</sup> |

|  |  |
| --- | --- |
| Total | 1302 |

- 1) Cultivars in plain text are standard cultivars designated to each HTR in the MAFF guideline (MAFF,1998). - 2) Cultivars in bold text are recently released cultivars registered in the RCC database with their HTR classified following the MAFF guideline. - 3) HTR based on NARO (2018) - 4) HTR based on NARO (2011) - 5) HTR based on Wada et al. (2010). - 6) HTR based on Mizobuchi et al. (2018)

Of the 1297 observations, 245 contained CG by type, such as milky-white, basal-white, belly-white, or back-white grains. This subset of data, referred to as “CG-type subset data”, inclluded 30 cultivars covering 5 HTRs and was used to examine the environmental responses of each CG type separately.

Weather data were extracted from the gridded daily meteorological dataset with a 1-km resolution developed by NARO (Ishigooka et al.,2017). We averaged daily mean temperature, solar radiation (SR), and relative humidity (RH) for 20 days after heading, a critical period affecting the occurrence of CG (Morita,2008). Where shading treatments were imposed, the incident SR was corrected using the shading ratio reported in each study. We derived TaHD as a heat stress index for rice quality, defined as the cumulative mean air temperature above the threshold temperature (T_b_) for 20 days after heading, following earlier research (Usui et al.,2016; Ishigooka et al.,2017; Nishimori et al.,2020) as follows;

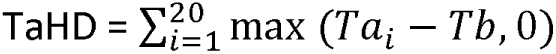

Ta_*i*_ is the daily mean temperature on the i-th day from the heading date (Table 1). Tb was set at 26 °C in our study based on a preliminary test that showed that 26 °C was appropriate across different HTRs (Supplementary Figure 1), in agreement with previous studies (Usui et al.,2016; Ishigooka et al.,2017; Nishimori et al.,2020).

### 2.3 Statistical analysis

All analyses were conducted using R (R Core Team,2022). We fitted a parametric linear mixed effects (LME) model and a non-parametric machine learning model (random forest, RF) to the whole dataset od 1297 observations to quantify the effects of weather variables (TaHD, mean RH, and SR) and HTR on CG. The following logit transformation was applied to stabilize the variance of CG proportion data (Yandell,1997), logit (CG) = ln (CG/(100-CG)). The variables TaHD, mean RH, and SR were treated as numerical variables and HTR as a factor. The use of two very different types of models (LME and RF) allowed us to test the robustness of the results to the model assumptions.

The following LME model was applied to examine the effects of HTRs and environmental factors on CG;

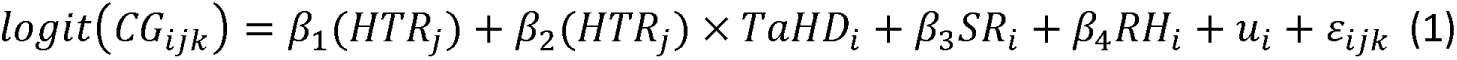

where i, j, and k are the indices of the reference-site (i.e., unique combination of article reference and site name), cultivar, and sub-treatment number for each site-cultivar, {3_l_,…, {3_4_ are fixed parameters, *u*_*i*_ is a random reference-site effect, *c*_*ijk*_ is a random residual term. The sub-treatment includes different nitrogen applications, planting patterns, or previous crops. Only TaHD × HTR interaction was included in this model because our preliminary tests indicated interactions between other environmental variables and HTR were not significant. Another preliminary test revealed that the inclusion of soil classification (group) did not improve the model goodness of fit, and so it was not included in the model. The model was fitted using the lme4 R package (Bates et al.,2015). The significance of the effects of explanatory variables was tested using ANOVA, and differences among HTRs were tested using the marginal means of CG estimated using the emmeans package (Lenth,2022). We also used the metafor package (Viechtbauer,2010) to illustrate CG responses to climatic factors with 95 % confidence intervals computed using bootstrapping.

We also used an RF model to determine the effects of HTR and climatic factors on CG without making a linear assumption and without the need to select any specific interaction (all interactions are taken into account in RF). This model was implemented using the randomForest package with the following settings (mtry = 2, ntree = 500) (Liaw and Wiener, 2001). CG was also logit-transformed before training RF. The importance of variables in the RF model was evaluated with the mean increase in mean square error (MSE) (computed by random permutation of the model input values), and the partial dependence plot function was used as an analysis tool to detect influential explanatory variables and then show how the average RF model responses are influenced by each predictor.

Both LME and RF models were fitted/trained with 80% of the total observations selected randomly while keeping the same proportions of HTR categories as in the entire dataset using the “sample” function in R. The fitted/trained models were then evaluated by computing the root mean square error (RMSE) with the test data, which accounted for 20% of all observations.

To examine the effects of HTR and environmental factors on different types of CG, the same LME model (Eq. (1)) was fitted to CG by type recorded in the sub-datasets, leading to three separate models. CG data by type were logit-transformed, but data with CG = 0% (n = 5) were excluded because they could not be logit-transformed. As a result, a subset of 240 observations was used for this analysis. For each CG type, an LME model was trained using 80% of all observations randomly selected from the subset, following the method described earlier. The root mean square error (RMSE) was calculated using 20% of the remaining data. We also fitted an LME model to the whole CG data (sum of three types) in the same manner, and we compared the model accuracy of CG estimated in two ways to determine whether the predictions of CG by type were more accurate than those obtained from a single model fitted to the aggregated CG.

## 3. Results

### 3.1 Grain quality observed in rice cultivars with different heat tolerance

The dataset used in our analysis contained 1297 CG observations for 48 cultivars of five different HTRs, obtained at 44 sites (196 site × year combinations) across Japan. HTR5 (intermediate tolerance level) recorded the most observations, accounting for 41% of the dataset, followed by HTR3 (susceptible, 23%) (Table 2). HTR4 (moderately susceptible) had the fewest observations but still had over 80 observations. CG spanned from 0 to 100 % (Supplementary Figure 2a), with a skewed distribution to the right. The overall mean CG was 20.7 %, 5.4 % points(percentage-points) greater than the median (15.3 %) (Table 3). Logit-transformed CG data, however, showed a bell-shaped symmetric distribution (Supplementary Figure 2b). The average values of CG obtained for each HTR class ranged from 12 to 29%, and declined as HTR increased (Table 3).

**Table 3.**
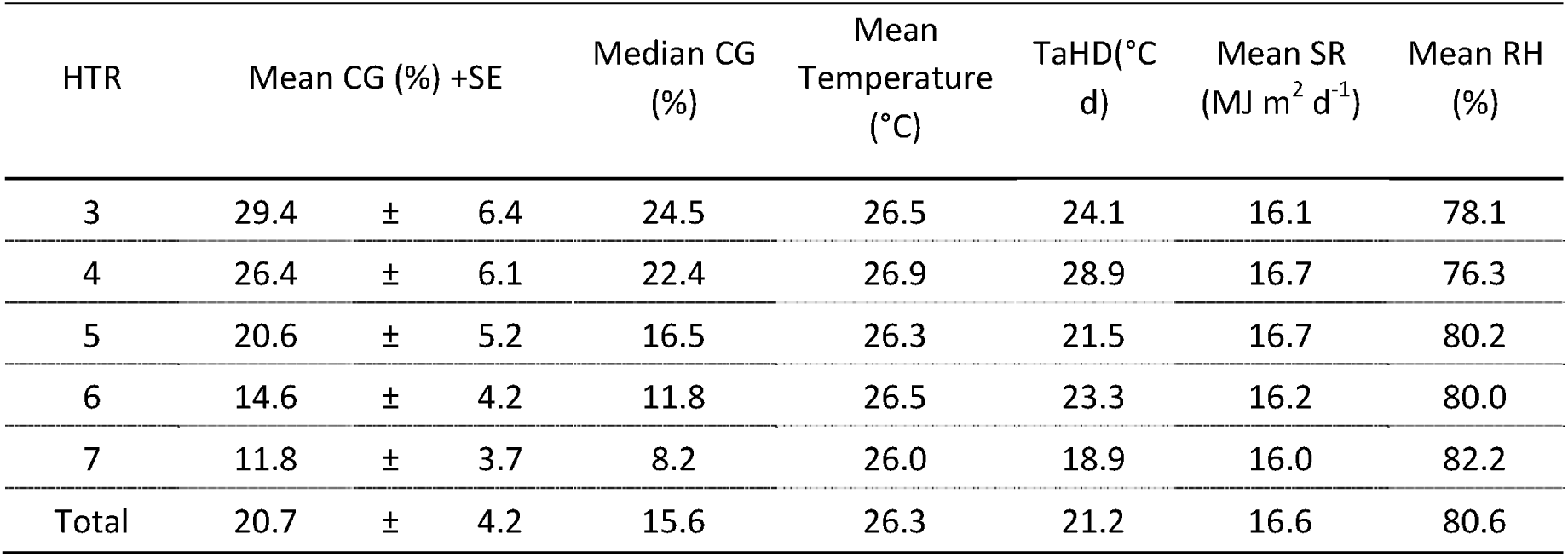
Mean measured CG (%), mean and standard error and weather conditions (averaged over 20 days after heading) for each HTR.

The logit-transformed CG data also exhibited a symmetric bell-shaped distribution when analyzed for each type of CG separately (Supplementary Fig. 4). The mean CG obtained for each HTR ranged from 9 to 28%, similar to the whole dataset. All three CG types (milky-white, basal-white, and belly/back white) decreased with increasing HTR (Supplementary Figure 5).

Air temperature, relative humidity (RH), and solar radiation (SR) averaged over 20 days after heading varied greatly among 196 site-years. The mean air temperature ranged from 20.8 °C to 30 °C with a median of 26.5 °C, implying that more than half of the observations were exposed to temperatures above the threshold temperature of 26°C (Supplementary Figure 2c). Mean SR and RH ranged from 7.4 to 23.8 MJ m^-2^ d^-1^ and from 64.2 % to 94.7 %, respectively (Supplementary Figures 2d and 2e). Weather conditions were similar among HTRs (Table 3, Supplementary Figure 2c-e). CG increased almost linearly with TaHD in all HTRs, but the increase in CG values was greater in low HTR levels than in high HTR levels (Figure 3).

**Figure 3.**
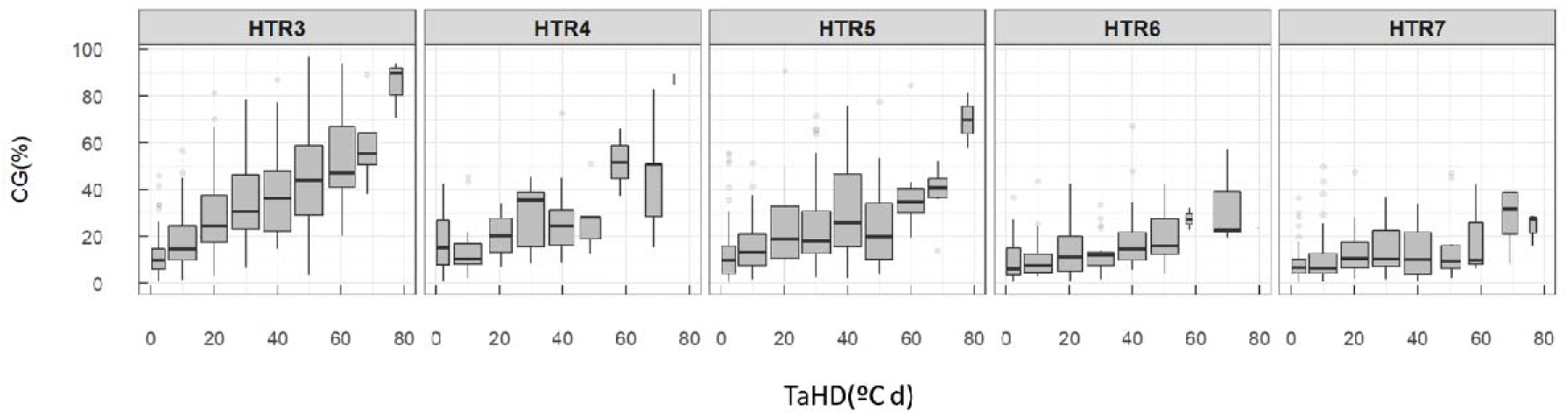
Boxplot of measured CG (%) in relationship to TaHD (°C d, representing the sum of temperature above 26°C, 20 days after heading) for each HTR category. Each box indicates the interquartile range (IQR) and the middle line in the box represents the median. The upper- and lower-end of whiskers are median 1.5lll×lllIQRlll±lllmedian. Open circles are values outside the 1.5lll×lllIQR.

### 3.2 Model comparison

We first applied the LME model (Eq.1) to test if the effect of temperature on CG differed among HTRs and if other environmental factors had a significant effect on CG. Overall, the model explained 63% of the CG when accounting for the reference-site effects, and 33% without (i.e., based on climate- and HTR-related inputs only). The RMSE of LME with the test data set was about 14% without accounting for reference-site effects (Figure 4).

**Figure 4.**
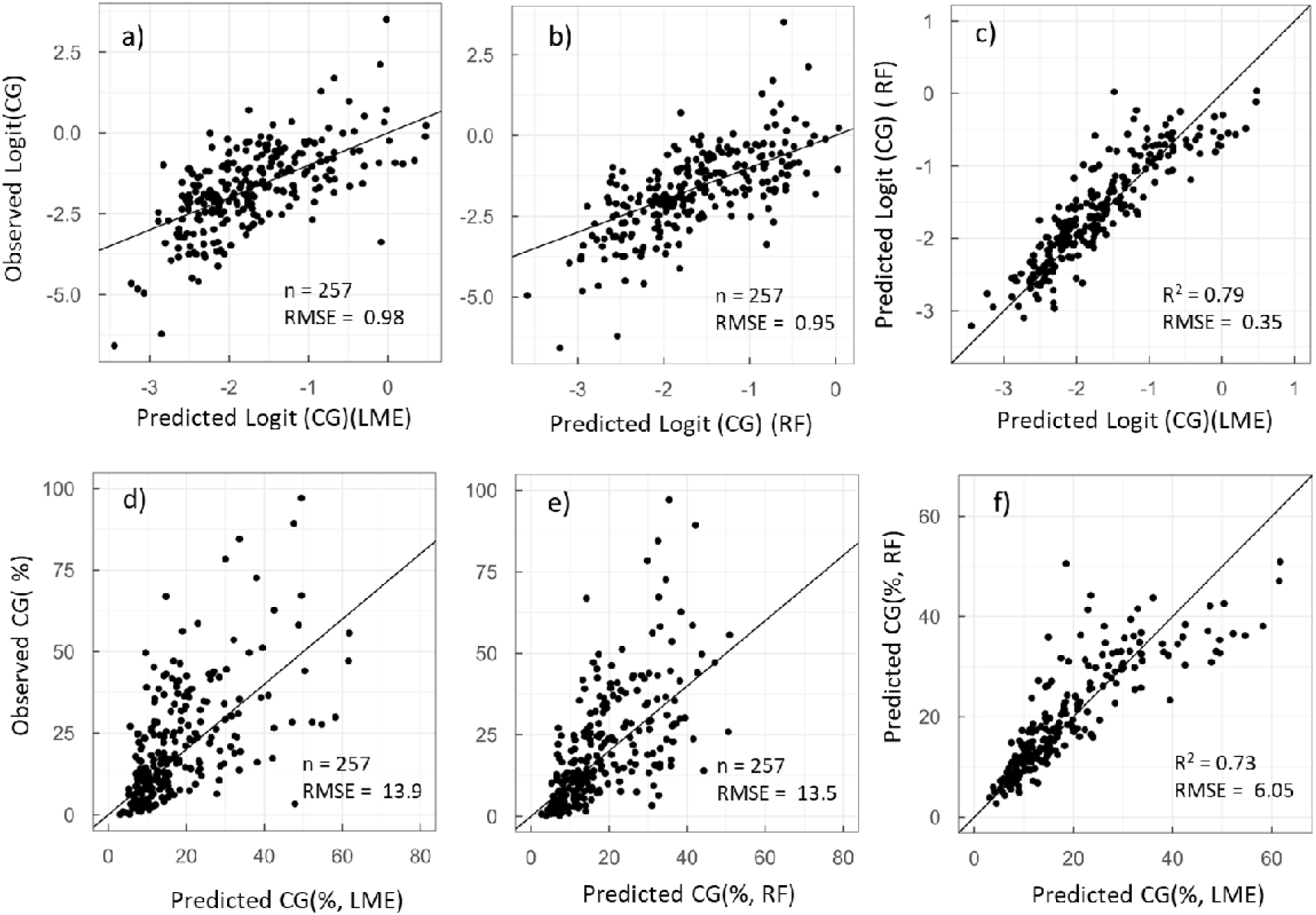
Relationships between observed and predicted values from the LME (a in logit scale, d in %) and the RF model (b in logit scale, e in %), and between the predicted values of the two models (c in logit scale, f in %).

The LME results revealed a strong effect of TaHD on CG (Table 4, P < 0.001), and a significant interaction between TaHD and HTRs (P < 0.001). The TaHD effect (i.e. coefficient) was greatest in HTR3 and decreased progressively as HTRs increased. Intercepts also differed significantly among HTRs, being smaller in HTR5, 6, and 7 than in HTR3 (P < 0.001). The effects of SR and RH were both significantly positive (P < 0.005 and P < 0.001, respectively), with RH having a larger influence than SR (Table 4). The marginal mean CG values revealed that increasing HTRs from 4 to 5 could significantly reduce CG by 7.7 % points and that increasing HTRs from 5 to 6 would decrease CG even further, by 5.0 % points, when the other environmental factors (SR and RH) were set to their mean values.

**Table 4.** ANOVA results of the rice LME model, estimated parameters, and estimated marginal means for each HTR.

|  | Variables | Mean Square | DFn<sup>(1)</sup> | DFd<sup>(2)</sup> | F value | p<sup>(3)</sup> |
| --- | --- | --- | --- | --- | --- | --- |
| ANOVA | TaHD | 105.6 | 1 | 1117 | 176.1 | 0.000*** |
|  | HTR | 15.06 | 4 | 1257 | 24.99 | 0.000*** |
|  | SR | 4.534 | 1 | 811 | 7.558 | 0.006** |
|  | RH | 25.70 | 1 | 1137 | 42.85 | 0.000*** |
|  | TaHD x HTR | 4.325 | 4 | 1236 | 7.209 | 0.000*** |
|  | HTR | Estimate<sup>(4)</sup> | SE | df | t value | P |
| Intercepts |  |  |  |  |  |  |
|  | HTR3 | -7.314 | 0.855 | 987 | -8.557 | 0.000*** |
|  | HTR4-HTR3 | 0.025 | 0.193 | 1282 | 0.127 | 0.899 |
|  | HTR5-HTR3 | -0.501 | 0.102 | 1270 | -4.922 | 0.000*** |
|  | HTR6-HTR3 | -0.906 | 0.131 | 1248 | -6.891 | 0.000*** |
|  | HTR7-HTR3 | -1.020 | 0.114 | 1242 | -8.933 | 0.000*** |
| Coefficients for covariates |  |  |  |  |  |  |
| TaHD | HTR3 | 0.046 | 0.003 | 1245 | 13.56 | 0.000*** |
|  | HTR4-HTR3 | -0.012 | 0.005 | 1242 | -2.204 | 0.028* |
|  | HTR5-HTR3 | -0.011 | 0.003 | 1245 | -3.351 | 0.001** |
|  | HTR6-HTR3 | -0.014 | 0.004 | 1228 | -3.352 | 0.001** |
|  | HTR7-HTR3 | -0.020 | 0.004 | 1227 | -5.156 | 0.000*** |
| SR |  | 0.042 | 0.015 | 811 | 2.749 | 0.006** |
| RH |  | 0.057 | 0.009 | 1137 | 6.546 | 0.000*** |
|  |  | Estimated marginal mean CG (%)<sup>(5)</sup> | Confidence limits (%) |  |  |  |
|  |  |  | Lower | Upper |  |  |
|  | HTR 3 | 26.5 a | 22.8 | 30.8 |  |  |
|  | HTR 4 | 22.3 a | 17.8 | 27.3 |  |  |
|  | HTR 5 | 14.6 b | 12.3 | 17.2 |  |  |
|  | HTR 6 | 9.6 c | 7.9 | 11.7 |  |  |
|  | HTR 7 | 7.7 d | 6.2 | 9.3 |  |  |

1) Degree of freedom for the numerator
2) Degree of freedom for the denominator estimated using the Kenward-Roger method in the lme4 R package (Bates et al., 2015).
3) \*\*\*, P < 0.001; \*\*, P < 0.01; and \*, P < 0.05 .
4) Estimated parameters in Eq. (1) based on the logit-transformed CG.
5) Estimated marginal means based on the untransformed CG. Values followed by the same alphabetic letters are not statistically different at P=0.05 using Tukey's method. Note that the difference between HTR6 and 7 was found significant, whereas that between HTR 3 and 4 was not, despite it being greater in the latter comparison. This was because the pairwise comparison was made using the logit-transformed data; the difference was greater in the HTR6-7 comparison than in the HTR3-4.

We also tested the RF model using the same explanatory variables and found that it explained 49% of the CG variation of the train data set. The RMSE with the test data set was about 14%, comparable to that of LME (Figure 4). The variable importance analysis of the RF model showed that TaHD was the most influential variable, followed by HTR, RH, and SR, in close agreement with the LME model (Table 4, Supplementary figure 3).

The LME model applied to the CG by type showed that all three CG types increased with TaHD, but that the TaHD effect tended to become progressively smaller with increasing HTRs, as evidenced by the significant interaction between HTR and TaHD (Supplementary Table 2). The effect of SR was not significant, while that of RH was significant except for basal-white (P<0.001 for milky-white and belly/back-white). Still, the estimates of coefficients were all positive in agreement with those for the whole CG (Supplementary Figure 6). We compared the accuracy of CG prediction obtained from the model fitted for the aggregated CG with that from the models fitted separately for three CG types and found that both predictions gave similar RMSE (15.4 % and 16.5 %, respectively) (Supplementary Figure 7).

### 3.3 Responses of CG to temperature, solar radiation, and humidity

Both models predicted similar responses of Logit (CG) to TaHD, SR, and RH (Supplementary Figure 3). The partial dependence plots of the RF model indicated that Logit(CG) responded almost linearly to all climatic variables studied, confirming that the linear assumption of the model LME was plausible. Since the RF and LME models showed similar goodness of fit (Section 3.2) and responses to climatic factors, we chose the LME model to describe the responses of CG to TaHD, RH, and SR for the different HTRs, due to its simplicity and interpretability.

CG increased as a function of TaHD for all HTRs, but the response was much weaker for high HTRs than for low HTRs (Figure 5). When SR and RH were fixed to their mean observed values, increasing TaHD from 20 to 80 °C d was projected to increase CG by 66 % points for HTR3, 45 % points for HTR5, and 19 % points for HTR7, confirming the effectiveness of heat-tolerant cultivars in reducing the negative impacts of high temperatures. The projected responses also showed that CG increased with both increasing RH and SR (Figure 5) to a greater extent for low HTRs than for high HTRs. When TaHD and SR were held constant, increasing RH from 70 to 90% was projected to increase CG by 22 % points for HTR3, 14 % points for HTR5, and 8 % points for HTR7. The effect of HTR was smaller for SR; increasing SR from 10 to 22 MJ m was estimated to raise CG by 9.8 % points for HTR3, 6 % points for HTR5, and only 4 % points for HTR7 (Figure 5)

**Figure 5.**
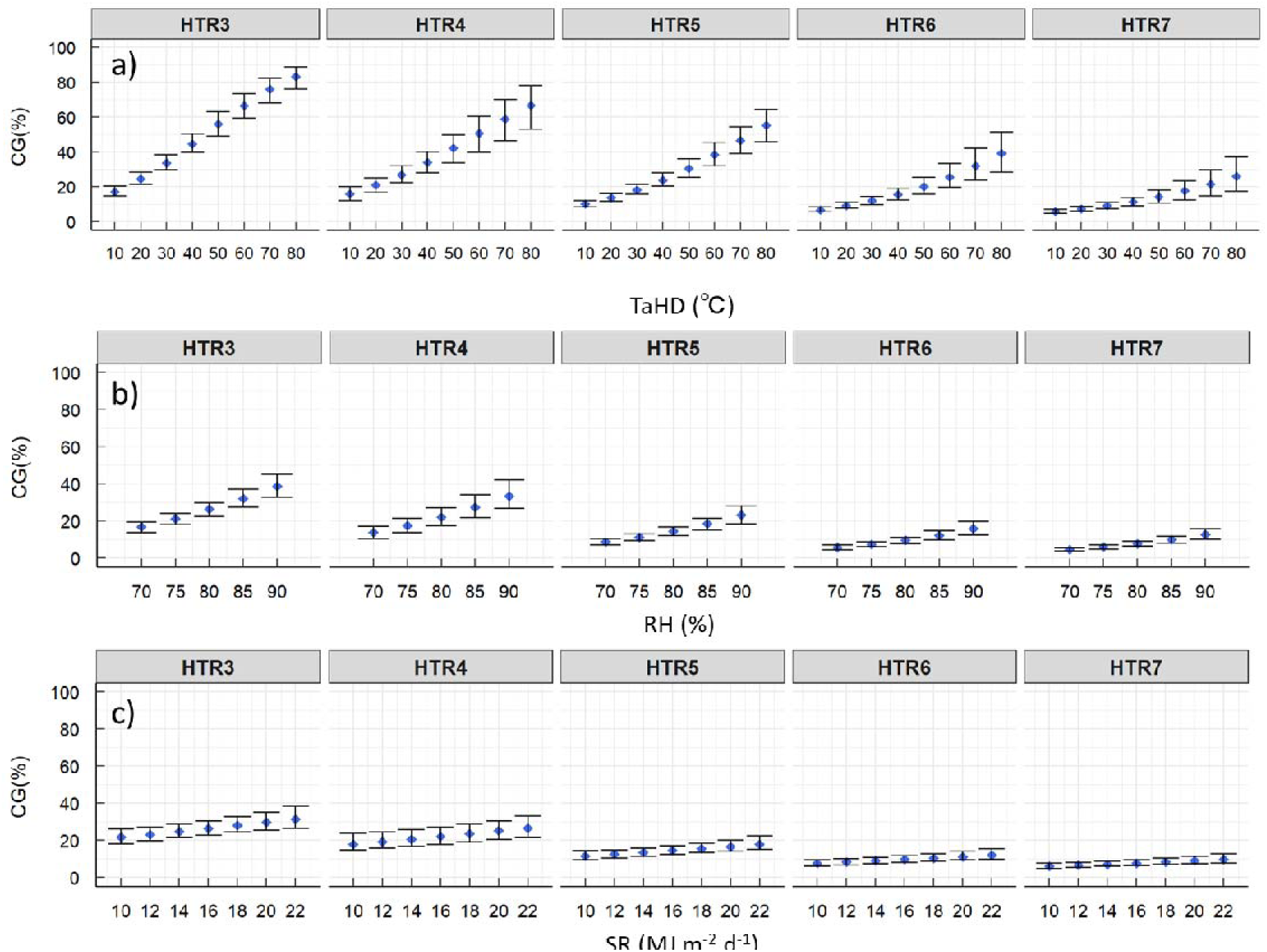
Response of CG (%) to TaHD (a), RH (b), and SR (c) estimated with the LME model for each heat tolerance rank (HTR). Error bars show 95% confidence intervals. Response to each weather variable was drawn by fixing other two environmental variables at their average values in the dataset: SR = 16 MJ m^-2^ d^-1^, RH = 80%, and TaHD = 22 °C d.

## 4. Discussion

Using a systematic literature search strategy, we collected 2,332 field trials of 321 cultivars conducted at 46 experimental stations across Japan to develop a quantitative effectiveness measure against heat damage on rice quality. The dataset covers all regions except Hokkaido, where temperatures are low and heat damage has yet to be reported. The dataset contains established national-level standardized cultivars with contrasting levels of heat tolerance. All the data were obtained in open fields, without additional warming, in conditions close to real farms. The dataset also includes key climatic variables and geographical information for each observation, facilitating the analysis of quality-climate relationships.

We tested two different models; a parametric model based on mixed-effect regression (LME) and a non-parametric model based on the random forest (RF) algorithm. The LME model assumes a linear relationship between the response and explanatory variables, while the RF model makes no linear assumption but is more difficult to interpret (Xie et al.,2021). Our results demonstrated that the two models agreed on their capacity to predict the CG variation in the test data (RMSE = 14 %), the importance of explanatory variables (TaHD > HTR > RH > SR), and the linear responses of CG to weather-related variables. Both models were able to detect the negative effect of high temperature on quality, which was found to be highly dependent on cultivars’ HTR. Each HTR group includes 6–15 cultivars (Table 2), bred in different locations and years from various backgrounds. They were selected as reference cultivars for each HTR based on their performance under high grain-filling temperatures (mean temperatures for 20 days after heading are typically around 27–28 °C). (Kaji et al.,2016; Tamura et al.,2018). The proposed models demonstrated that increasing HTR could reduce the occurrence of CG at high temperatures (>26 °C), and we found that a shift of HTR by one rank could decrease CG by more than 5%.

Grading of rice grain quality in Japan is determined based on several criteria, the most important one being the percentage of undamaged grains as described in Article 3 of the Agricultural Products Inspection Law, where the first-grade rice must have at least 70% of undamaged grains. Chalky grains are considered immature and one of the major causes of lowering the rice grade, particularly under high temperatures during the grain filling period. Earlier, Nishimori et al. (2020) showed that, from a nationwide survey in Japan, the percentage of first-grade rice sharply dropped by 3-4 % points each % point rise in CG when CG exceeded 5 %. When CG exceeded 25 %, the first-grade rice % was nearly zero and was replaced by second or third-grade rice. Reducing CG by raising HTR can help to maintain appearance quality (Table 4) and, thus, quality grade in hotter climates. For instance, at a mean grain-filling temperature of 27 °C (0.6 °C higher than the average of the entire dataset, Table 3, equivalent to TaHD = 20 °C d), CG was estimated to be 24.5 % (95% CI ranging from 21.2 to 27.75%) for HTR3 (susceptible) (Figure 5), implying that it is already difficult for HRT3 cultivars to produce first-grade rice. At the same grain-filling temperature, CG was calculated to be 13.6 % (95% CI, 11.4 ∼ 15.9 %) with HTR5 (intermediate) and 9.0 % (7.5∼10.9) with HTR6 (moderately tolerant) (Figure 5), indicating that raising HTR by just one step could substantially contribute to attaining first-grade rice. When the air temperature further rises, however, the adaptation limit is reached, even with HTR6 or 7. For instance, at a grain-filling temperature of 28.5 °C (TaHD = 50 °C d), CG was projected to reach 20% (16.2∼25.4) for HTR6 and 14% (11.1∼18.1) for HTR7, indicating that the reduction of first-grade rice is inevitable, calling for further improvement in heat tolerance.

Our results confirmed that grain-filling temperature is the dominant climatic factor affecting CG. The variable TaHD (cumulative temperature above the threshold value 26°C) was found to be a key factor, as reported elsewhere (Nishimori et al.,2020). Some researchers used a different base temperature for different genotypes, assuming that heat tolerance would raise the threshold temperature (Takimoto et al.,2019; Masutomi et al.,2022). We also examined this possibility by varying the base temperature for each HTR, but found that the best model fit was obtained with a base temperature of about 26 °C regardless of HTRs (Supplementary Figure 1). We, therefore, fixed the base temperature at 26 °C for all HTRs. It is possible that the intercepts of the LME model, which also differed significantly among HTRs, accounted for the CG variation near the threshold temperature.

Two other climatic factors, Relative Humidity (RH) and Solar Radiation (SR), were also found to have significant effects on CG according to the LME model. CG increased with both RH and SR, but the effect of RH was stronger, with the F value being 43 for RH and 7.6 for SR (Table 4). To our knowledge, only a few have studies reported the effects of RH on CG; one of them used a growth chamber to impose RH treatments, and the other statistically analyzed climatic factors on CG of field-grown rice in wet and dry seasons (Wakamatsu et al.,2009; Zhao and Fitzgerald,2013), both demonstrating that CG increased with RH in line with the current findings. The effect of SR on CG has been frequently reported (Deng et al.,2018; Ishimaru et al.,2018), with evidence that a specific type of chalk, milky-white grains, which depends on assimilate supply (Kobata et al.,2004; Tsukaguchi and Iida,2008), increased under low SR (Ishimaru et al.,2018). Some modelling studies have already considered the effects of SR (Okada et al.,2011; Yoshida et al.,2016; Takimoto et al.,2019), accounting for the rise in CG when solar radiation is limited. Our analysis showed, however, that high SR had a positive impact on CG although the effect of SR was much smaller than that of TaHD or RH (Table 4, Figure 5). Our additional analysis based on a subset data for three types of CG revealed that the dependence of CG on SR was either positive or close to 0 in all CG types (Supplementary Table2 and Figure 6). This could be explained in part by the fact that an important mechanism by which SR increases CG despite its positive effects on assimilate supply is greater incident radiation, which raises panicle temperatures. High RH could also raise panicle temperatures by reducing transpirational cooling (Yoshimoto et al.,2022). The fact that both RH and SR increased CG suggests that the indirect impacts of these climatic factors on CG via panicle temperatures override a beneficial effect of SR through increased assimilate supply, which could reduce the occurrence of milky-white grains. It is important to note that increases in CG with rising RH and SR decreased with higher HTRs, implying that the effects of RH and SR were heat-mediated (Figure 5).

Our analysis by CG type revealed that HTR was effective in reducing all three types of CG (milky-white, basal-white, and belly/back-white grains). The analysis also confirmed the significant TaHD x HTR interactions, with a stronger interaction for basal-white and belly/back-white grains (Supplementary Table 2 and Figure 6) for their sensitivity to high temperatures (Wakamatsu et al.,2007). Our model comparison did not show any advantage in distinguishing different types of CG, and thus supports the use of the whole CG as the response variable (Supplementary Figure 7).

Our models can play at least two roles in genetic improvements in heat tolerance. First, when combined with climate scenarios, the model can be used to backcast quantitative breeding targets that are temporally and spatially explicit. Breeding targets can also be defined based on global warming levels. Because the lead time in breeding is often long, the breeding target—heat tolerance level in the current context—must be developed well before variety release. Current temperatures during the grain-filling period vary across regions. For instance, our dataset showed grain-filling temperatures averaged for each prefecture in the recent decade, ranged from below 25 °C to above 28 °C. They are projected to rise over time as global warming proceeds, strongly suggesting that the breeding target for HTR needs to be tailored for each region and time. The current models can estimate CG from major climatic factors for different tolerance levels and can help design locally specific targets. The models are built on the Japanese rice dataset but might be applied to other countries/regions where standard cultivars are already established. Second, the models may be used to derive objective and quantitative heat tolerance ranks for new cultivars based on field trials and their environmental conditions. Heat tolerance ranks of new cultivars are currently determined based on relative cultivar performances, established comparing candidate lines with standard cultivars. The proposed models may potentially assist expert judgment by providing quantitative tolerance levels from multiple field data sets, although further testing of the model is needed for this purpose. Heat tolerance is required for other characteristics, particularly heat-induced spikelet sterility, which can substantially affect cereal yield (Ye et al.,2021). However, because the growth processes and temperature sensitivity for spikelet sterility are different from those for grain quality in threshold values and sensitive stages (Sanchez et al.,2014), modelling and enhancing heat tolerance for high-temperature-induced spikelet sterility requires different strategies that should be addressed separately from the present study.

In principle, the proposed models could be used to estimate the effects of weather- and cultivar-related inputs, but many other factors are involved in the CG variation. The LME model explained 63 % of the variations when accounting for the reference-site effects. Other factors may include some management factors such as nitrogen application (Wakamatsu et al.,2008; Tanaka et al.,2010), planting density (Morita et al.,2012), and water management (Chiba et al.,2017). In this study, most data were from cultivar trials on experimental stations where standard levels of fertilizers were applied and sufficient irrigation was possible, which might have decreased the CG variation compared to farmers’ fields. Experimental sites had different soils, but adding soil groups did not improve the prediction accuracy (data not shown). Nevertheless, diverse fertilizer applications, soil fertility, and irrigation methods could exist in actual farmers’ fields. Model adjustments, incorporating more covariates related to soil or management practices, may be necessary for the model to be widely applicable in the future.

Because the model is empirical, it cannot be extrapolated. The maximum TaHD in the dataset was 80 °C d, which is equivalent to the average grain-filling temperature of 30 °C. According to the climate projections based on various greenhouse gas emissions, the mean air temperature in Japan may rise by 0.6 ∼ 3.0 °C and by 1.1 ∼ 6.7 °C from the 1981-2000 period by mid- and late-century, respectively (Ishigooka et al.,2017), suggesting that grain-filling temperatures could exceed 30 °C, particularly in currently warm regions. In addition, the current model does not account for atmospheric CO_2_ concentration or wind speed. Notably, the effect of CO_2_ is relevant in the context of climate change because elevated CO_2_ exacerbates the heat-induced degradation of rice appearance quality (Yang et al.,2007; Usui et al.,2016), which could dramatically influence the projected impacts of climate change on grain appearance quality (Ishigooka et al.,2021). There are a few reports that show heat-tolerant cultivars effectively sustain appearance quality under elevated CO_2_ levels (Usui et al.,2014; Hasegawa et al.,2019), but this needs to be examined more extensively. Further expansion of the rice CG dataset and model improvement will be needed.

## Conclusions

We aimed to quantify the effects of improving cultivars’ heat tolerance ranks (HTRs) to alleviate heat damage on rice grain quality using recently established national-level standardized cultivar heat tolerance ranks in Japan. A linear mixed effect model and a random forest model were developed and tested with a dataset containing 1,297 field trials of 48 cultivars from 44 experimental stations across Japan, covering all regions where high temperatures are a source of concerns. The two models agreed on their capacity to predict the variation in chalky grain percentage (CG) in test data (RMSE = 14 or 15%), the importance of explanatory variables, and linear responses to climatic variables. The proposed models demonstrated that increasing HTR could reduce the occurrence of CG, especially under high temperature conditions. Increasing the HTR by a single step (from intermediate to moderately tolerant) can significantly increase the chances of obtaining first-grade quality rice under grain filling temperatures of up to 27°C, but tolerance levels must be improved if more extreme temperatures are expected. The proposed models can be used to backcast quantitative, temporally, and spatially explicit breeding targets under climate change.

## Acknowledgements

This study was partly supported by the Environment Research and Technology Development Fund (JPMEERF20S11820) of the Environmental Restoration and Conservation Agency of Japan; JST Grant Number JPMJPF2013 supported this work; the Joint Linkage Call supported by NARO and INRAE; and the project CLIMAE of INRAE.

