## Supplementary figures and tables for "Effectiveness of heat tolerance rice cultivars in preserving grain appearance quality under high temperatures - A meta-analysis"

**Corresponding Autor**

Toshihiro Hasegawa

**Institute for Agro-Environmental Sciences**

**National Agriculture and Food Research Organization (NARO)**

**3-1-3 Kannondai, Tsukuba, Ibaraki 305-8604, Japan**

|  | Cultivars |
| --- | --- |
| Standard  Cultivars registered by  MAFF | Akanezora, Akisayaka, Aoinokaze, Hatsuboshi, Hinohikari, Komanomai, Matsuribare, Sainokagayaki, Sasanishiki, Satojiman, Tomohonami, Kinuhikari, Koganebare, Sinrei, Tachiharuka, Akitakomachi, Haenuki, Hitomebore, Koshihikari, Mutsuhomare, Nikomaru, Nipponbare, Mizuhonokagayaki, Akisakari, Fukei227, Hanaechizen, Koganemasari, Kokoromachi, Mineharuka, Natsuhonoka, Nishihikari, Satonouta, Shifukunominori, Tochiginohoshi, Tsuyahime, Natsuhonoka, Eminokizuna, Fusaotome, Otentosodachi |
| Other Heat-tolerant cultivars | Akihonami, Fufufu, Harumoni, Koisomeshi, Yumiazusa, Chihominori, Inahokkori, Koinoyokan, Akiharuka, Kumasannnochikara, Koshijiwase, Nijinokirameki, Tsukiakari, Toyama81, Tsuyakirari Emidawara, Kankinokaze, Genkitsukushi, Natsuhikari, Mie23, Oidemai, Sagabiyori, Sainokizuna, Yosakoibijin |

**Supplementary Table1** List of cultivars used for the systematic literature search

**Supplementary: Optimum base temperature (Tb) for TaHD calculation**

To determine the optimum base temperature (Tb) for the cumulative mean air temperature for 20 days after heading (TaHD) that explains well the response of CG to heat stress, we tested the LME model against the CG dataset by changing Tb from 24 to 28°C. TaHD was derived as follows:

TaHD = $\sum_{i=1}^{20} max({Ta}_{i}-Tb,0)$

where Ta_i_ is the daily mean temperature on the ith day after the heading date.

Firstly, we used the following LME model for each HTR:

$logit\left( {CG}_{ik} \right)=\beta_{0}+\beta_{1}{TaHD}_{i}+\beta_{2}{SR}_{i}+\beta_{3}{RH}_{i}+u_{i}+\varepsilon_{ik}$	(Eq.1)

where i is the reference-site index (in combination), k is the sub-treatment number for each site and cultivar, β0～3 are fixed parameters, u_i_ is a random reference-site effect, and ε_ik_ is a random residual term. Fitting Eq.1 was repeated for TaHD calculated with Tb varied from 24 to 28°C in a 1°C step.

Then the following LME model including HTR and its interaction with TaHD was fitted to the CG dataset:

$logit\left( {CG}_{ijk} \right)=\beta_{0}+\beta_{1}({HTR}_{j})+\beta_{2}({HTR}_{j})\times{TaHD}_{i}+\beta_{3}{SR}_{i}+\beta_{4}{RH}_{i}+u_{i}+\varepsilon_{ijk}$ (Eq.2)

where i, j, and k are the reference-site (in combination), cultivar, and sub-treatment number indices for each site-cultivar, 04 are fixed parameters, ui is a random reference-site effect, and ijk is a random residual term. Fitting Eq.2 was also repeated using various TaHD obtained from various Tbs.

Additionally, we used average air temperature (Tave) for 20 days after heading instead of TaHD to confirm the advantage of using TaHD instead of Tave.

As shown in Supplemental Figure 1, the AIC with Tave was substantially greater than that with TaHD estimated for almost all Tb values examined, showing the advantage of utilizing TaHD over Tave. The AIC of the LME model was lowest at Tb, around 26 °C, across all HTRs, with no clear evidence of Tb increasing with HTR. The AIC from Equation 2 using the entire dataset confirms that TaHD calculated with Tb = 26 °C provides the best fit (Supplementary Figure f). We, therefore, used the fixed Tb value of 26 °C for all HTRs.

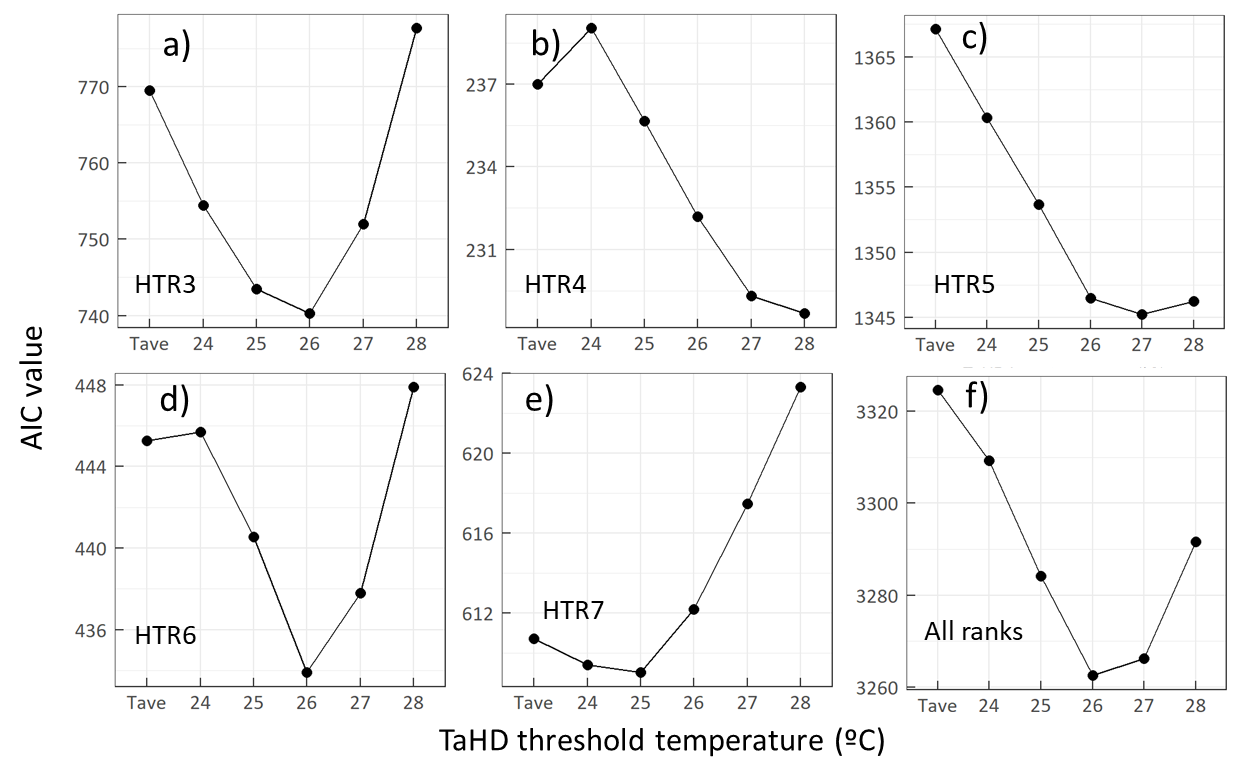

**Supplementary Figure 1.** Changes in AIC values of the LME models based on TaHD with Tb, for each HTR (a to e) and the whole dataset with all HTRs (f). Tave is for the model using average air temperature instead of TaHD.

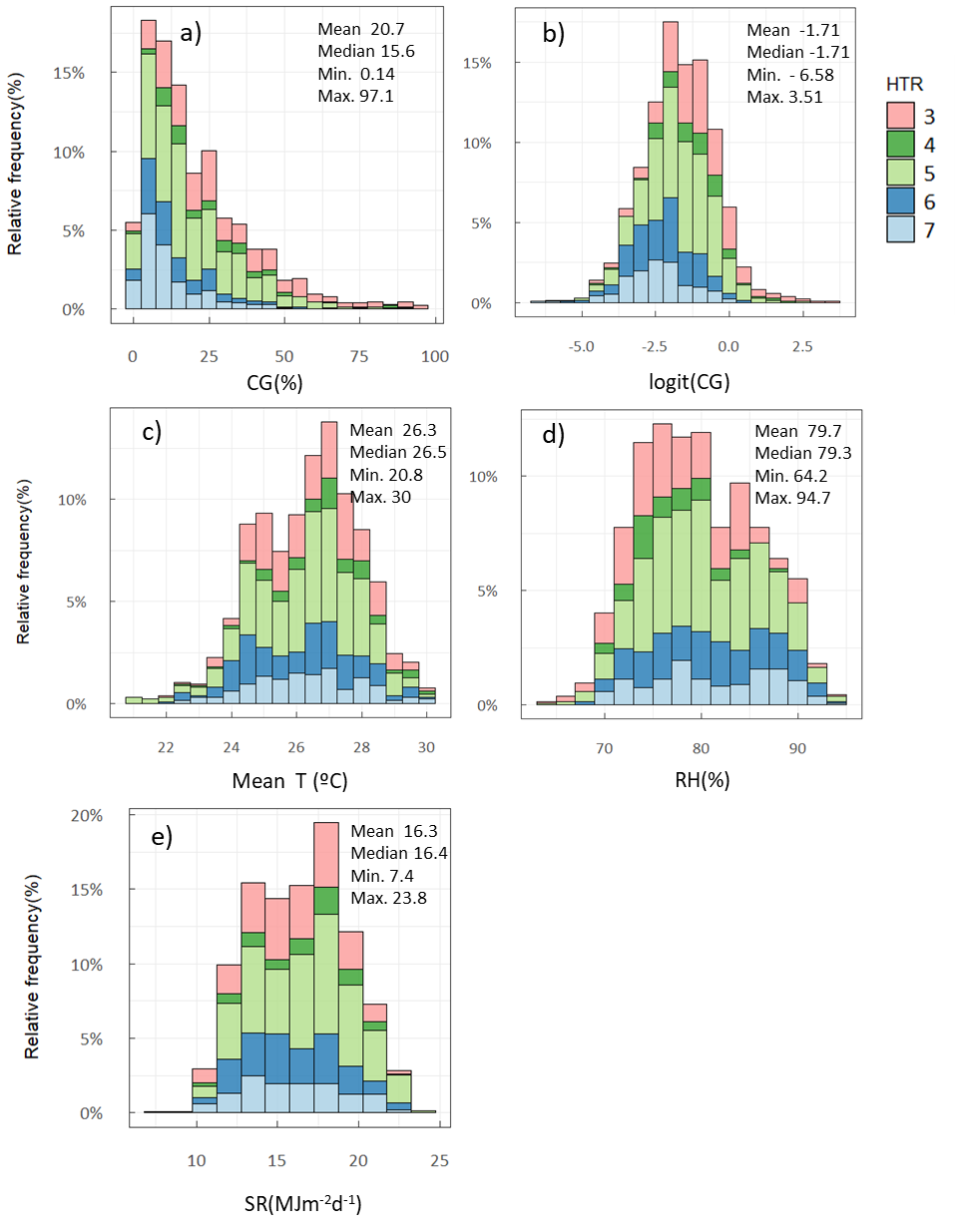

**Supplementary Figure 2.** Relative frequency of main variables in the dataset. (a) CG (%), (b) logit-transformed CG, (c) mean air temperature (°C), (d) relative humidity (RH, %), and e) solar radiation (SR, MJ m^-2^d^-1^). All weather factors are averages for 20 days after heading. n=1302.

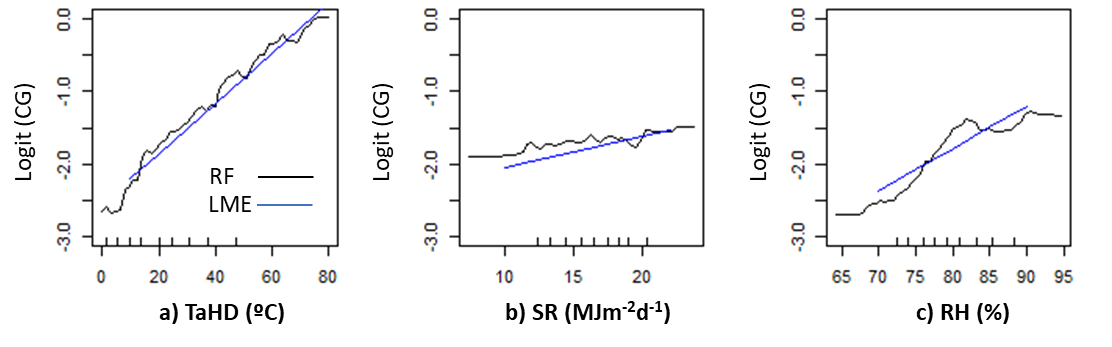

**Supplementary Figure 3.** Partial dependence plot of logit(CG) showing the average responses to a) TaHD (°Cd), b) SR (MJm^-2^d^-1^), c) RH (%) simulated by the RF model, and the linear relationships between logit (CG) and each weather variable (a-c) fitted by the LME model for HTR5.

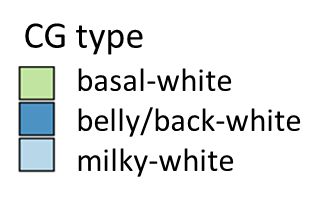

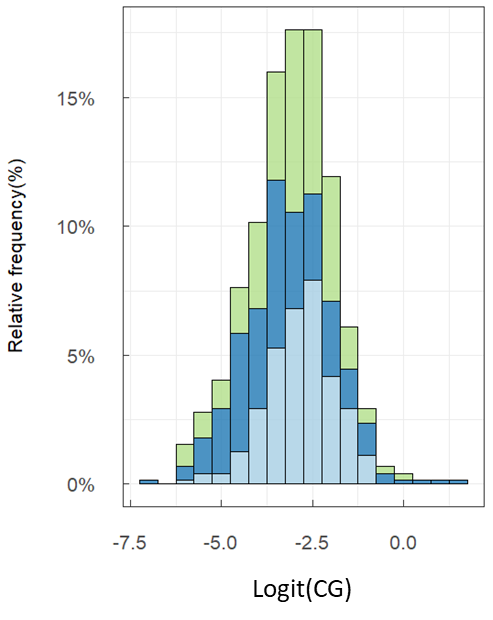

**Supplementary Figure 4.** Relative frequency of logit-transformed CG by types (basal-white, belly/back white, milky-white) in the subset of data reporting CG types (n = 240).

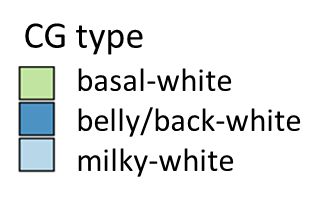

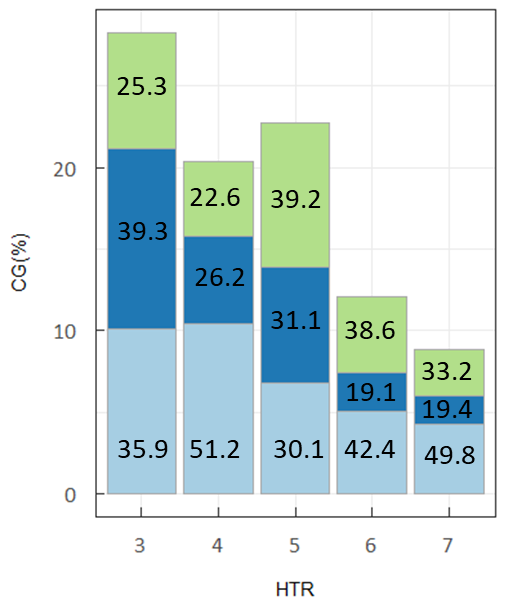

**Supplementary Figure 5.** Values of CG (total and per CG type) by Heat Tolerance Rank (HTR).

Numbers inside each stack bar indicate the percentage of each CG type (basal-white, belly/back white, milky-white) in the subset of data reporting CG types (n = 240).

**Supplementary Table2.** ANOVA results of the LME model fitted for each CG type (n = 240)

| CG type | Variables | Mean Sq | DFn^(1)^ | DFd^(2)^ | F value | *P*^(3)^ | |
| --- | --- | --- | --- | --- | --- | --- | --- |
| milky-white | TaHD | 2.84 | 1 | 223.63 | 6.14 | 0.014 | * |
|  | HTR | 2.41 | 4 | 219.83 | 5.21 | 0.001 | *** |
|  | SR | 0.32 | 1 | 147.61 | 0.69 | 0.406 |  |
|  | RH | 4.96 | 1 | 156.02 | 10.70 | 0.001 | ** |
|  | TaHD x HTR | 1.00 | 4 | 212.85 | 2.16 | 0.075 | . |
| basal-white | TaHD | 7.24 | 1 | 226.46 | 11.89 | 0.001 | *** |
|  | HTR | 2.49 | 4 | 214.76 | 4.08 | 0.003 | ** |
|  | SR | 0.00 | 1 | 148.15 | 0.00 | 0.982 | ns |
|  | RH | 0.47 | 1 | 160.62 | 0.78 | 0.379 | ns |
|  | TaHD x HTR | 2.06 | 4 | 204.95 | 3.38 | 0.010 | * |
| belly/back-white | TaHD | 14.84 | 1 | 226.58 | 20.24 | 0.000 | *** |
|  | HTR | 3.18 | 4 | 218.28 | 4.34 | 0.002 | ** |
|  | SR | 1.81 | 1 | 161.03 | 2.47 | 0.118 | ns |
|  | RH | 8.04 | 1 | 171.81 | 10.98 | 0.001 | ** |
|  | TaHD x HTR | 2.65 | 4 | 210.75 | 3.62 | 0.007 | ** |

1) Degree of freedom for the numerator

2) Degree of freedom for the denominator estimated using the Kenward-Roger method in the lme4 R package (Bates et al., 2015).

3) ***, P< 0.001; **, P < 0.01; and *, P<0.05

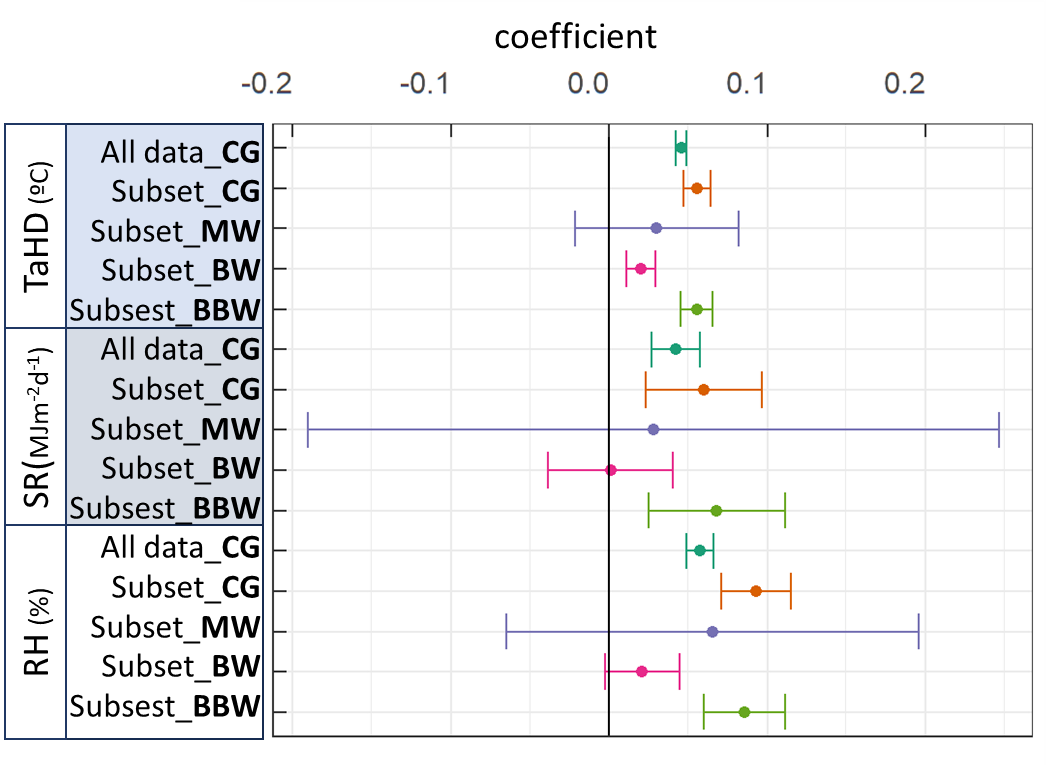

**Supplementary Figure 6.** Comparison of the dependence of chalky grains on meteorological variables derived from a LME model fitted to the whole dataset (n=1297), from LME models fitted to the three CG-types separately (n=240), and from a LME model fitted to all CG types taken together but with the subset of data for which the CG types were reported (“Subset CG", n=240). Points indicate the estimated regression coefficients for TaHD, SR, and RH obtained with the LME models. Error bars are the standard errors of each estimated coefficient. Bold letters denote each response variable; MW stands for “milky-white grain”, BW stands for “Basal-white grain”, and BBW stands for “Belly/back-white grain”.

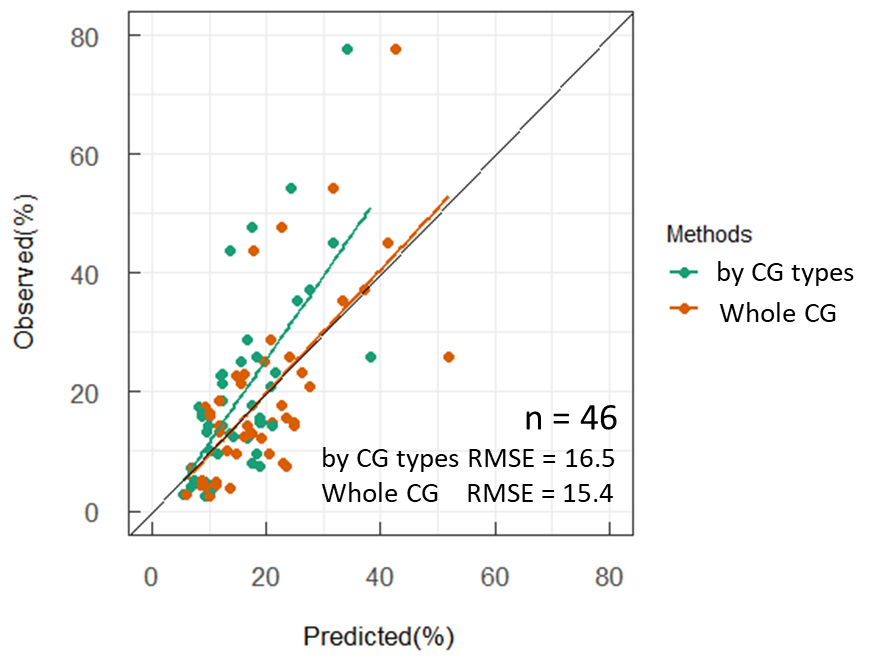

**Supplementary Figure 7.** Relationship between observed and predicted values from two types of LME models tested against data not used for parameterization (n = 46). The green dots are derived from LME models for three different types of chalky grains separately, where values predicted for each type were summed to estimate the total CG. The orange dots were derived from the LME model fitted against the total CG.

**Reference list identified from the search on four science research databases (the second source)**

1. Arakawa, M., Ooka, N., Minoda, T., Ishii, H., Takei, Y., Ueno, T., Tokura, K., Saito, K., Shigematsu, O., Kozasu, M., Okada, Y., Watanabe, K., Arai, M., Arai, N., Noda, S., 2009. Breeding of a new rice cultivar "Sainohohoemi" with disease and pest combined resistance, excellent palatability for double cropping. Bulletin of the Saitama Prefectural Agriculture and Forestry Research Center 9, 21-34.
2. Asami, H., Miura, Y., Watanabe, T., Uno, T., Tajima, R., Saito, M., Ito, T., 2020. Silicate fertilizer improves grain quality in brewers’ rice (*Oryza sativa* L.) ‘Toyonishiki’. Japanese Journal of Soil Science and Plant Nutrition 91, 11-20.
3. Fujii, Y., Hatakeyama, S., Mitsukawa, M., Matsuno, H., 2004. Factors related to white core rice of cultivar "Hinohikari" occurrence in the normal season culture, 1997. Kyusyu Agricultural research 66, 11.
4. Hamachi, Y., Miyazaki, M., Tsubone, M., Oono, Y., Odahara, K., 2012. Effects of High Air Temperature in the Summer of 2010 on the Kernel Quality of the Heat-tolerant Rice Cultivar “Genkitsukushi”. Japanese Journal of Crop Science 81, 332-338.
5. Hamagashira, A., Ide, Y., Sugiura, K., Nakamura, M., Tsuda, K., Kato, M., Ikeda, A., Sugiura, N., Ito, A., Matsumoto, Y., Mizukami, Y., Mori, K.-i., Watanabe, Y., Ando, Y., Takikawa, T., Shimada, T., Nakajima, Y., Nezu, T., 2020. Breeding of a new rice variety "Aichi 135" with tolerance to high temperature during maturation Research Bulletin of Aichi Agricultural Research Center 52, 31-39.
6. Hasegawa, W., Shiraishi, M., Onari, S., Yasui, T., Emoto, K., Sato, Y., Kitazono, K., Jufuku, K., Yamasaki, A., Nagamoto, Y., Ono, K., Ninomiya, Y., Goto, S., Kawazu, K., 2009. Characteristics of rice varieties "Nikomaru" and "Akimasari" urgently introduced for global warming in Oita prefecture. Bulletin of Oita Prefectural Agriculture, Forestry and Fisheries Research Center, Agriculture Section 3, 27-44.
7. He, B., Toyata, M., Kusutani, A., 2017. Effects of Sowing Date on the Growth, Yield and Palatability of Japanese Rice Cultivated by Non-tillage Direct Sowing in Kagawa Prefecture. Japanese Journal of Crop Science 86, 267-275.
8. Hiroichi, S., Sasaki, S., Watanabe, Y., Kuchiki, Y., Saito, T., Kobayashi, N., Sato, M., 2017. Varietal characteristics of a New Rice Cultivar “Satoyamanotubu” in Fukushima Prefecture. Tohoku Journal of Crop Science 60, 1-4.
9. Ito, A., Funao, T., Shirotani, M., Kato, M., Sugiura, K., Nakamura, M., Kato, T., 2012. The selection of standard rice varieties under high-temperature grain filling conditions in extremely early season culture in Aichi prefecture. Research Bulletin of Aichi Agricultural Research Center 44, 45-51.
10. Kakiuchi, J., Kawanishi, T., Morimoto, T., 2008. Analytical Studies of the Factors Affecting the Quality of Rice Cultivar ‘Kinuhikari' in Wakayama (2) Effect of Midseason Drainage, Planting Density, and method of fertilizer application on the yield and quality. Bulletin of the Wakayama Research Center of Agriculture, Forestry and Fisheries 9, 1-13.
11. Kimura, H., 2018. A study of cultivation conditions for rice cultivar "Nikomaru" to be ranked special A Bulletin of the Ehime Research Insutitute of Agriculture, Forestry and Fisheries 10, 1-9.
12. Kinoda, N., Sakaiya, E., Oyamada, Z., Anamizu, K., 1995. Harvest time and grain quality of rice cultivar "Mutsuhomare" and "Maihime" ripened under the hot weather in 1994. Tohoku agricultural research 48, 23-24.
13. Kinoshita, N., Mitsukawa, M., Fujii, Y., 2019. "Kumasannnokagayaki", a new rice cultivar with excellent eating quality, tolerance to high temperature during maturing period and good grain appearance Agriculture and horticulture 94, 113-121.
14. Kobayashi, A., Sugimoto, K., Hayashi, T., Kondo, M., Sonoda, J., Tukaguchi, T., Wada, T., Yamanouchi, U., Iwasawa, N., Yano, M., Tomita, K., 2016. Development of a near isogenic line of ‘Koshihikari’ with a seed dormancy gene and an evaluation of its resistance to heat-induced quality decline. Breeding Research 18, 1-10.
15. Kojima, Y., Ebitani, T., Omoteno, M., Kidani, Y., Yamaguchi, T., Mukaino, N., Kaneda, H., Takarada, T., Doi, M., Fukuda, M., Ishibashi, T., 2008. New rice cultivar "Tenkomori". Bulletin of the Toyama Agricultural Research Center 25.
16. Komaki, Y., Sasahara, H., Uehara, Y., 2002. Varietal Differences of Ripening Ability among Early-maturing Rice Varieties under High Temperature in a Vinyl House. The Hokuriku Crop Science 37, 12-16.
17. Maeda, S., Senoo, T., Sugimoto, S., 2015. Estimation of suitable area and time of transplanting based on weather conditions for rice cultivar "Kinumusume", using rice growth model in Okayama prefecture. Kinki Chugoku Shikoku agricultural research 26, 23-32.
18. Maeda, S., Watanabe, T., 2013. Grain filling characteristics of "Nikomaru", a Rice cultivar with tolerance to high-temperature during ripening period. Bulletin of the Research Institute for Agriculture Okayama prefectural technology center for Agriculture, Forestry, and Fisheries 4, 1-8.
19. Miyai, R., Kawamura, K., Adachi, Y., 2015. Effects of transplanting season, fertilizer and harvesting time, yield and grain quality in paddy rice variety "Kinumusume". Bulletin of the Wakayama Research Center of Agriculture, Forestry and Fisheries 3, 19-26.
20. Miyazaki, M., Uchikawa, O., Tanaka, K., Fukushima, Y., 2008. The grain quality of rice and the rice plant nutrition under the different transplanting time. Report of Kyusyu Branch Crop Science Society of Japan 74, 11-13.
21. Miyazaki, M., Yoshino, M., Uchikawa, O., Iwabuchi, T., Araki, M., Ishitsuka, A., Odahara, K., 2011. Effects of different transplanting periods and nitrogen applications on the yield per unit, grain quality, and eating quality of a new rice cultivar "Genkitsukushi". Bulletin of Fukuoka Agricultural Research Center 30, 18-24.
22. Morita, K., Matsushima, T., Yamaguchi, T., Saito, A., Furuhata, M., 2012. Evaluation of the Dense Planting in Late Transplanting Aimed to Avoid High Temperatures During the Ripening Period in Rice Cultivar Koshihikari. Japanese Journal of Crop Science 81, 349-356.
23. Morita, K., Takahashi, W., Sugimori, F., Furuhata, M., 2011. Planting Density Suitable for Late Transplanting for Avoiding High Temperature During the Ripening Period of Rice Cultivar Koshihikari in Toyama Prefecture, Japan. Japanese Journal of Crop Science 80, 220-228.
24. Morita, S., Iwabuchi, T., Makiyama, S., Koga, J., Tanaka, Y., Kira, T., Yabuoshi, Y., Wakamatsu, K.-i., Kureya, K., Wakiyama, Y., Sakai, M., Wada, T., Hirota, Y., Fujii, Y., Hasegawa, W., Nagayoshi, Y., Komaki, Y., Simoyama, N., Tanaka, K., Kitagawa, H., Hruguchi, S., Mitsukawa, M., Tsuji, T., Asakawa, M., 2010. Analysis of the Actual Situation and Factors behind the Recent Decline in Paddy Rice Production and Quality in Kyushu and Future Responses. Memoirs of the National Agricultural Research Center for Kyusyu Okinawa Region 94, 1-105.
25. Mukoyama, T., Shimura, T., Ishii, T., Watanabe, A., Ueno, N., 2019. Cultivation characteristics, quality and ripening ability under high temperature of "Tsuyahime", a medium rice variety with good palatability, in Yamanashi prefecture. Bulletin of the Yamanashi Prefectural Agricultural Technology Center 11, 1-8.
26. Nagahata, H., Kuroda, A., 2004. The Effects of High Temperature Treatment on the Occurrence of White-immature Kernels and the Trait of Eating Quality among Early-maturing Varieties of Rice. The Hokuriku Crop Science 39, 81-84.
27. Nagano, K., Chiba, B., Sasaki, K., Endo, T., Wagatuma, K., Hayasaka, H., 2017. A new rice cultivar "Tohoku 194". Bulletin of the Miyagi Prefectural Furukawa Agricultural Eperiment station 12, 1-18.
28. Nakagawa, J., Mori, S., 2012. Selection of Standard Rice Cultivars for Ripening Capability under High Temperature Conditions in Shiga Prefecture. Journal of Crop Research 57, 23-31.
29. Nakagawa, J., Yoshida, T., Mori, S., Hino, K., Yamada, Y., Miyamura, H., Nishitani, K., 2014. A new rice cultivar "Mizukagami" with tolerance to high temperature during ripening stage. Bulletin of the Shiga Agricultural Technology Promotion Center 52, 1-14.
30. Ogawa, M., HIrooka, M., Okubo, E., Mori, Y., 2020. Optimum fertilization and Appropriate harvest timing for paddy rice cultivar "Inahokkori". Bulletin of Gunma Agricultural Technology Center 17, 27-34.
31. Okada, Y., Ishi, H., 2017. Development of high-temperature injury mitigation technique for "Sainokagayaki". Bulletin of the Saitama Agricultural Technology Research Center 16, 15-32.
32. Oshima, Y., 2009. Cultivation characteristics and method of fertilizer application for the paddy rice "Satojiman". Bulletin of the Kanagawa Agricultural Technology Center 151, 51-56.
33. Oshima, Y., 2009. Influence of poor sunshine at the middle stage, high temperature at the initial stage and high temperature with dryness at the maturity stage of grain growth on the quality of brown rice. Bulletin of the Kanagawa Agricultural Technology Center 151, 39-50.
34. Sasahara, H., Goto, A., Shigemune, A., Nagaoka, I., Komaki, Y., Ymaguchi, M., Maeda, H., Matsushita, K., Miura, K., 2018. A new variety for sushi "Eminokizuna". Bulletin of the NARO, Agricultural Research for Central Region 5, 1-18.
35. Sasaki, K., 2017. Analysis of rice variety with good taste on premature transplant in Yamagata. Tohoku agricultural research 70, 15-16.
36. Sasaki, R., Nakai, J., Fujita, M., Kosaka, Y., Matsumoto, J.-i., Ueda, N., Adachi, Y., Kadowaki, S., Tsukimori, H., Watanabe, T., Katsuba, Z., Mkatsukasa, M., Yamamoto, Z., Fujita, K., Taniguchi, H., Takata, S., Sawada, T., Matsumoto, S., Ishii, T., Iwai, M., Senoo, T., Ymaguchi, K., Ikegami, M., Okubo, K., Ishii, T., Nagata, K., 2012. A comprehensive review on the impact of recent high temerature on rice grain ripenning in western region of Japan, and the adaptaion and mitigation technologies. Miscellaneous publication of NARO Western Region Agricultural Research Center 9, 41-146.
37. Sato, H., Sasaki, S., Otera, M., Kikuchi, N., 2018. An evaluation of grain quality by rice appearance analyzer in brewers' rice. Tohoku Journal of Crop Science 61, 5-8.
38. Takamatsu, M., 2018. Breeding of Kazesayaka and direction of breeding. The Hokuriku Crop Science 53, 57-58.
39. Takata, S., Sakata, M., Kameshima, M., Yamamoto, Y., Miyazaki, A., 2010. Evaluation Method of Ripening Capability of Rice Varieties under High Temperature and Poor Sunshine Conditions in Early-Season Culture in Warm South-Western District of Japan. Japanese Journal of Crop Science 79, 142-149.
40. Takata, S., Sakata, M., Kameshima, M., Yamamoto, Y., Miyazaki, A., 2010. Varietal Difference in the Relation between the Occurrence of White Immature Kernels Caused by a High Temperature during the Ripening Period and the Amount of Basal Nitrogen Application in Rice. Japanese Journal of Crop Science 79, 150-157.
41. Tamura, K., Kataoka, T., Nakanishi, A., Sato, H., Tamura, Y., Sakai, M., Yamaguchi, O., Wada, T., Tsubone, M., Tatara, I., Tokuda, S., Yoshida, K., Koga, J., Nakayama, M., Fujii, Y., Mitsukawa, M., Shimizu, Y., Hasegawa, W., Shiraishi, M., Nagayoshi, Y., Matsuura, S., Wakamatsu, K.-i., Sato, M., Sonoda, J., Takeuchi, Y., 2018. Standard Rice Varieties with High Temperature Tolerance at the Ripening Stage in the Warm Region of Japan. Japanese Journal of Crop Science 87, 209-214.
42. Tanaka, K., Miyazaki, M., Uchikawa, O., Araki, M., 2010. Effects of the Nitrogen Nutrient Condition and Nitrogen Application on Kernel Quality of Rice. Japanese Journal of Crop Science 79, 450-459.
43. Usui, Y., Sakai, H., Tokida, T., Nakamura, H., Nakagawa, H., Hasegawa, T., 2014. Heat-tolerant rice cultivars retain grain appearance quality under free-air CO2 enrichment. Rice 7, 6.
44. Wakamatsu, K.-i., Sasaki, O., Uezono, I., Tanaka, A., 2007. Effects of High Air Temperature during the Ripening Period on the Grain Quality of Rice in Warm Regions of Japan. Japanese Journal of Crop Science 76, 71-78.
45. Yamaguchi, T., Kojima, Y., Ebitani, T., Kaneda, H., Kidani, Y., Doi, M., Ishibashi, T., Mukaino, N., Omoteno, M., Takarada, T., Yamamoto, Y., 2006. A new rice cultivar "Tentakaku" Bulletin of Toyama Agricultural Research Centre 23, 29-43.
46. Yamazaki, S., Yuzawa, M., Nagashima, H., Aonuma, S.-i., Miyoshi, M., Shinozaki, A., Izawa, Y., Yamaguchi, M., 2012. A new paddy rice cultivar "Tochigi-no-hoshi". Bulletin of Tochigi Agricultural experimental station 68, 1-13.
47. Yokoyama, K., Takatori, H., Fujii, H., Ando, T., Watanabe, K., 2000. Feature of occurrence of Milky white kernel in rice cultivar "Haenuki" in Shonai district of Yamagata prefecture in 1999, and its environmental and physiological factors Tohoku agricultural research 53, 25-26.
48. Yoshinaga, S., Nagata, K., Shiratsuchi, H., Fukuda, A., 2012. Characteristics of Grain Quality and the Causal Factors in Direct-seeded Rice (*Oryza stiva* L.) in Lowland Field in a Cooler Region of Japan. Japanese Journal of Crop Science 81, 432-440.
49. Yoshinaga, S., Shiratsuchi, H., Nagata, K., Fukuda, A., Nakabayashi, M., Yokoyama, H., Kimura, T., Hikage, K., Odanaka, A., Asano, M., Mikami, Y., Shimazu, H., Kikawa, H., Miura, C., Wakamatsu, K., Yamakawa, A., Inoue, Y., Asanome, N., Nakayama, Y., Shimamune, T., Suzuki, Y., Kida, Y., Sasaki, S., 2008. Grain Filling Properties and Characteristics of Yield and Grain Quality of Direct-seeded Rice (*Oryza stiva* L.) in Tohoku District. Bulletin of TOHOKU Agricultural Research Center 109, 41-82.
